# Effect of unequal vaccination coverage and migration on long-term pathogen evolution in a metapopulation

**DOI:** 10.1101/2022.12.16.520709

**Authors:** Alicia Walter, Sylvain Gandon, Sébastien Lion

## Abstract

Worldwide inequalities in vaccine availability are expected to affect the spread and spatial distribution of infectious diseases. It is unclear, however, how spatial variation in vaccination coverage can affect the long-term evolution of pathogens. Here we use an analytical model and numerical simulations to analyse the influence of different imperfect vaccines on the potential evolution of pathogen virulence in a two-population model where vaccination coverage varies between populations. We focus on four vaccines, with different modes of action on the life-cycle of a pathogen infecting two host populations coupled by migration. We show that, for vaccines that reduce infection risk or transmissibility, spatial heterogeneity has little effect on pathogen prevalence and host mortality, and no effect on the evolution of pathogen virulence. In contrast, vaccines that reduce pathogen virulence can select for more virulent pathogens and may lead to the coexistence of different pathogen strains, depending on the degree of spatial heterogeneity in the metapopulation. This heterogeneity is driven by two parameters: pathogen migration and the difference in the vaccination rate between the two populations. We show that vaccines that only reduce pathogen virulence select mainly for a single pathogen strategy in the long term while vaccines that reduce both transmission and virulence can favor the coexistence of two pathogen genotypes. We discuss the implications and potential extensions of our analysis.

## 1 Introduction

A major challenge when designing public health measures is to balance the often urgent need to manage epidemic and endemic infectious diseases with potential long-term evolutionary side-effects (Dieckmann et al., 2005; Day et al., 2020). Vaccines that reduce pathogen spread or harmfulness are a key tool for the management of infectious diseases in human and animal populations. At the same time, vaccination, like other public health measures, may trigger an evolutionary response that could lead to pathogen adaptation and the erosion of vaccine efficacy. Even though vaccination has been shown to be more durable than other therapeutic interventions (Kennedy and Read, 2017, 2018), the use of imperfect vaccines may promote the evolution of pathogen traits, such as virulence, transmission or immune escape (Read et al., 2015; Gandon and Day, 2008; McLeod and Gandon, 2022). Evidence of vaccine-driven pathogen evolution is mixed, with some vaccines causing little or no vaccine adaptation (such as anti-measles vaccine), and other vaccines having been tied with increases in pathogen virulence (such as the various vaccines against Marek’s disease in poultry, Gandon and Day (2008); Read et al. (2015); Day et al. (2022)). Previous theoretical studies have shed light on this dichotomy, showing that evolution of pathogen life-history traits in response to vaccination can be favoured when the vaccine is imperfect and acts by reducing the cost of virulence (Gandon et al., 2001, 2003; Read et al., 2015), compared to vaccines that solely target pathogen reproduction by decreasing the susceptibility and transmissibility of hosts. This pattern has also been shown to be broadly preserved in the presence of spatial structure (Zurita-Gutiérrez and Lion, 2015) or temporal heterogeneity (Walter and Lion, 2021).

However, human and animal populations are often characterised by heterogeneity in vaccination coverage and complex patterns of dispersal between local populations. For instance, in human populations, different regions, even geographically close, may have very different vaccination coverages, as exemplified by regional differences in vaccination against measles in the Netherlands (Knol et al., 2013) or the worldwide inequality of access to COVID-19 vaccines (Mathieu et al., 2021; Rydland et al., 2022). Furthermore, the globalization of travel and trade tends to increase the connections between local populations and to facilitate the dispersal of hosts and pathogens, with potentially worrying consequences for pathogen spread and evolution (Memish et al., 2015; Boots and Sasaki, 1999). Hence, host populations are best viewed as metapopulations of local patches with different epidemiological characteristics, coupled by host or pathogen migration. It is not clear how the interplay between migration and spatial heterogeneity in vaccine coverage will affect the epidemiology and evolution of pathogens.

As a first step towards answering this question, we analyse a theoretical model to understand the combined effects of migration and vaccine distribution on the long-term evolution of pathogen virulence in a metapopulation. We consider the simple case of a metapopulation of two host populations connected by pathogen dispersal. Each population is characterised by a distinct vaccination coverage, which may lead to different local optima for the pathogen. As such, the problem we investigate is reminiscent of classical local adaptation models that have shown that a metapopulation with migration between populations with different optima can select for generalist or specialist strategies depending on migration and the degree of similarity between the populations (Meszéna et al., 1997; Day, 2000; Ronce and Kirkpatrick, 2001; Débarre et al., 2013; Mirrahimi and Gandon, 2020). Here, we extend these models to an epidemiolgoical context and ask how these factors may affect the evolution of pathogen polymorphism and what are the implications of such polymorphism for the epidemiological and evolutionary management of diseases.

Following Gandon et al. (2001), we assume the vaccine is imperfect, and can have different modes of action on the pathogen’s life cycle. Moreover, we consider a heterogeneous vaccine distribution across the two populations, and explore the epidemiological and evolutionary consequences of different vaccine distributions, ranging from homogeneous distributions (i.e. equal distribution between the two populations) to fully heterogeneous distributions (i.e. one population is completely vaccinated and the other completely naive). We investigate, both analytically and numerically, the long-term effects of spatial heterogeneity on the evolution of pathogen virulence, either for vaccines with a single mode of action, or for vaccines combining different modes of action on the pathogen’s lifecycle. We present and discuss the epidemiological and evolutionary consequences of these different scenarios, as well as the broader implications of our study for infectious disease management.

## 2 Epidemiological model

We consider a metapopulation composed of two populations *A* and *B*, coupled by parasite migration (life cycle depicted in **figure 1**, see table 1 for notations). In each population, the host-pathogen interaction is described by a Susceptible-Infected epidemiological model. Susceptible hosts are produced in each population at rate *b*, and vaccinated with a habitat-specific probability. In population *A*, a proportion *ν*_*A*_ of new hosts receive the vaccine and enter the vaccinated class 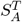, while a proportion 1 − *ν*_*A*_ enter the unvaccinated class 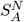 . Likewise, in population *B*, a proportion *ν*_*B*_ of new hosts enter the 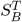 class, and a proportion 1 − *ν*_*B*_ enter the 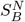 class. Note that this can be interpreted as a model of vaccination at birth, but we show in *Supplementary Information S*.*7* that an alternative model where adults are vaccinated leads to qualitatively identical results. All hosts die at a natural rate *d*. The metapopulation is infected by a pathogen that can migrate between the two populations at rate *m*, which varies between 0 and 0.5. For *m* = 0, there is only local transmission of pathogens and susceptible hosts of population *A* (respectively *B*) are only infected by infected hosts from population *A* (respectively *B*). For *m* = 0.5, susceptible hosts can be equally infected by hosts from population *A* or *B*. We consider that scenarios where *m* is greater than 0.5, meaning that susceptible hosts would be more infected by hosts from the other population, would be unrealistic (Park et al., 2015). Infected hosts in a given population transmit the pathogen at a rate that differs for naive (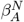 and 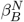) or vaccinated (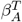 and 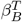) hosts. Similarly, infected hosts in population *j* and immunity state *k* have an additional disease induced mortality 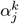. Note that we only consider pathogen migration, and, as a result, host densities are not directly altered by migration. A susceptible host can be infected either by a philopatric pathogen (with probability 1 − *m*) or by an allopatric pathogen (with probability *m*). This could correspond to at least three scenarios: (1) direct pathogen propagule dispersal, (2) environmental transmission through abiotic or biotic factors, or (3) rapid to-and-fro host movements between the two populations (e.g. with workers commuting between two cities). In addition, we assume for simplicity that migration is symmetric between the two populations.

**Table 1:** Table of parameters and variables.

| Parameter | Definition |
| --- | --- |
| Life history traits |  |
| $\alpha_j^k$ | disease-induced mortality caused by the pathogen strain in population $j$ and class $k$ |
| $\beta_j^k$ | transmission of the disease of the pathogen strain in population $j$ and class $k$ |
| $h_j$ | production of propagules by hosts in population $j$ |
| $H_j^k$ | force of infection caused by hosts in population $j$ on susceptible hosts in class $k$ |
| Reproductive values |  |
| $c_j$ | class reproductive value in population $j$ |
| Ecology |  |
| $b$ | influx of susceptible hosts |
| $d$ | natural death rate |
| $m$ | parasite migration rate |
| Vaccines |  |
| $\nu_j$ | probability of vaccinating new hosts in population $j$ |
| $\delta$ | $\nu_B - \nu_A$ |
| $r_1$ | efficacy of anti-infection vaccine |
| $r_2$ | efficacy of anti-growth vaccine |
| $r_3$ | efficacy of anti-transmission vaccine |
| $r_4$ | efficacy of anti-virulence vaccine |
| $\sigma$ | susceptibility of hosts ($1 - r_1$) |
| $k$ stands for naive ($N$) or vaccinated ($T$) hosts | |
| $j$ stands for population $A$ or $B$ | |

**Figure 1:**
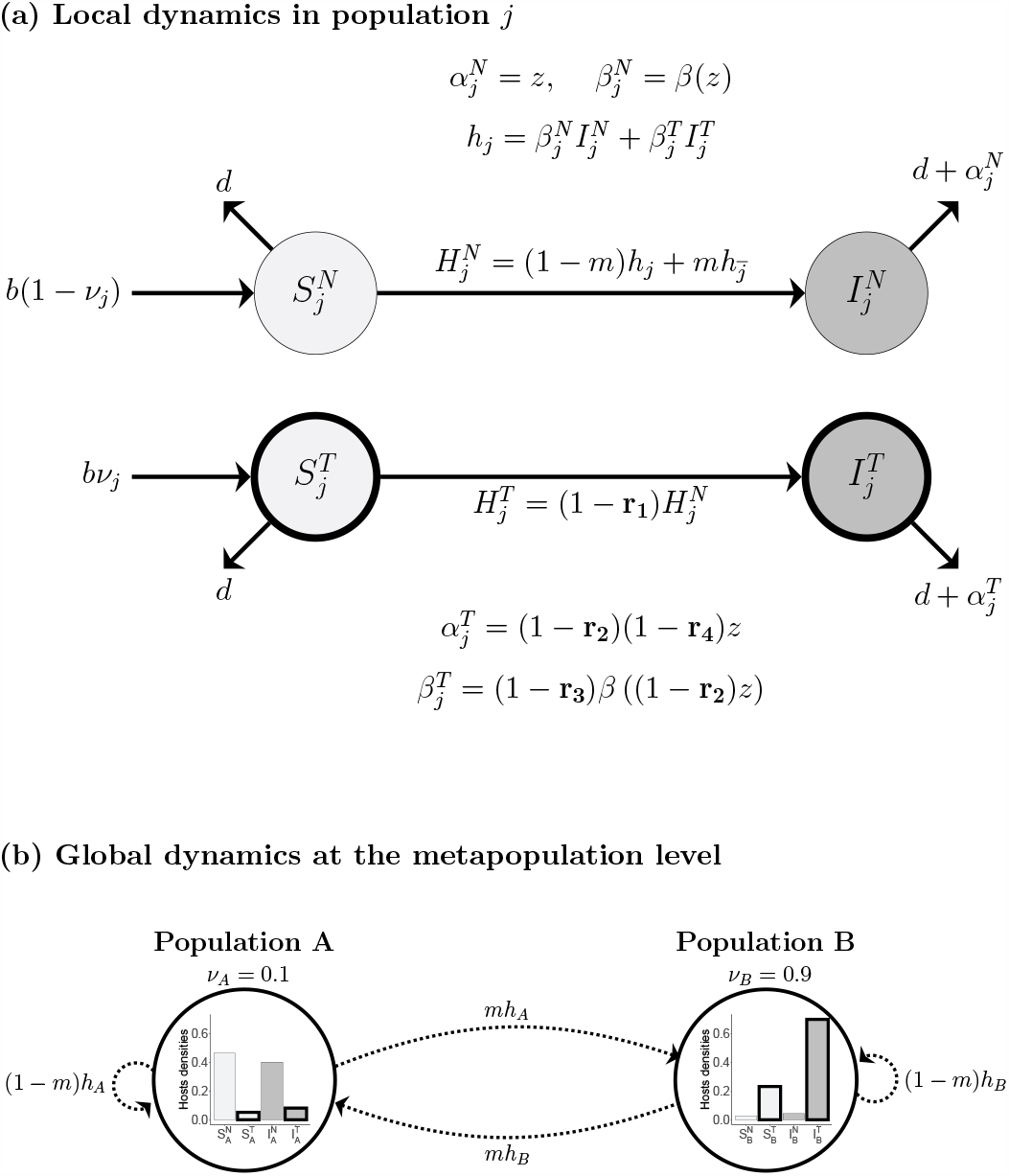
Schematic presentation of the model. (a) Life cycle of the host-pathogen interaction in the population *j*, showing transition rates between classes. Susceptible hosts are represented in light grey and infected hosts in dark grey, vaccinated hosts are in thick circle and naive hosts in thin. Four types of vaccines are considered: (1) anti-infection vaccines that reduce susceptibility to thep athogen (with efficacy *r*_1_), (2) anti-growth vaccines that reduce within-host replication and act on both transmission and virulence (with efficacy *r*_2_), (3) vaccines that only reduce transmissibility (with efficacty *r*_3_), and (4) vaccines that only reduce virulence (with efficacy *r*_4_). As shown in the figure, these vaccines affect the force of infection 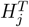, and the life history traits in the treated class (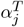 and 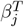). See the main text for more details about the mathematical expressions. (b) Transition rates in the metapopulation showing how pathogens migration *m* affects the forces of infection and the prevalence in each population. Barplots of hosts densities are shown as examples of populations structure at equilibrium with an anti-virulence vaccine, where *ν*_*A*_ = 0.1, *ν*_*B*_ = 0.9, *r*_4_ = 0.8, *m* = 0.01, *b* = 2 and *d* = 1.

These assumptions lead to the the following ODE system for the population *j* :

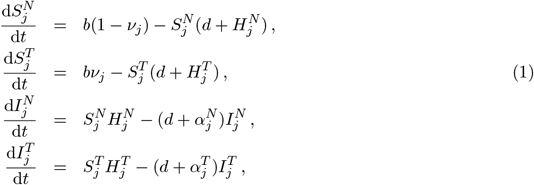

where *j* represents the population *A* or *B*, and 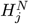 (resp. 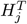) is the force of infection experienced by a naive (resp. vaccinated) susceptible host in population *j*. This force of infection depends on pathogen migration and is a weighted average of the contributions of infected hosts in populations *A* and *B*, so that 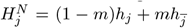, where we use the notation 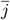 to refer to the non-focal population *j* (i.e., 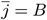 if *j* = *A*), and *h*_*j*_ is the force of infection due to hosts in population *j*:

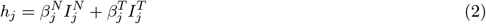

where 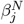 (resp. 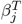) is the pathogen’s transmission rate in class *N* (resp. *T*) in population *j*. Furthermore, we assume that the force of infection on a vaccinated susceptible host in population 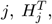, is reduced by a factor *σ* compared to that on a naive susceptible host, so that 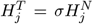 . The full 8-equation ODE system is given in *Supplementary Information S*.*1*.

Following Gandon et al. (2001, 2003), we consider four imperfect vaccines with different modes of actions on the pathogen’s lifecycle. We assume that the vaccine can either (1) reduce the susceptibility of hosts (anti-infection vaccine, noted *r*_1_), (2) reduce the within-host growth rate of pathogens (anti-growth vaccine, noted *r*_2_), (3) reduce pathogen transmission (anti-transmission vaccine, noted *r*_3_) and (4) reduce disease-induced mortality in infected hosts (anti-virulence vaccine, noted *r*_4_). Biological examples of each type of vaccine are presented in Walter and Lion (2021) and the relationship between the vaccine efficacy parameters *r*_*k*_ and the epidemiological parameters is given in figure 1a.

With these assumptions, the typical epidemiological behaviour of our model is either pathogen extinction or convergence to an endemic state where the densities of all classes of susceptible and infected hosts settle on equilibrium values. In the next section we use adaptive dynamics theory to analyse how the migration rate *m*, and the probabilities of vaccination *ν*_*A*_ and *ν*_*B*_ affect the long-term evolutionary outcome of the pathogen population, before applying the results to specific epidemiological scenarios.

## 3 Evolutionary dynamics

Assuming that the metapopulation infected by a resident strain of the pathogen has reached an epidemiological equilibrium, we consider a scenario where a mutant strain of the pathogen appears with different life-history traits, and investigate the invasion success of this mutant using the adaptive dynamics framework (Geritz et al., 1998; Dieckmann et al., 2005).

### 3.1 Mutant invasion fitness

Consider that the resident pathogen has trait *z* and that a mutant strain appears in the metapopulation with a different trait value, noted *z*^*′*^. We assume that both transmission and virulence are functions of the trait *z* (we will provide explicit examples for these functions in section 3.4) and, consequently, the mutation leads to different life-history traits for the mutant pathogen, denoted by a prime (e.g. 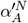 is the virulence caused by the mutant pathogen in naive infected hosts in population *A*). The mutant can either die out or invade and replace the resident strain. Mutant success depends on its invasion fitness, *R*(*z, z*^*′*^), that represents the number of surviving mutant offspring per mutant parent when the mutant allele is rare (Hurford et al., 2009; Otto and Day, 2011). Note, that this invasion fitness is measured per generation (i.e. it measures the expected number of secondary infections over the duration of a primary infection by the mutant) but an equivalent invasion fitness could be derived per unit of time. In our model this takes the form (*Supplementary Information S*.*2*) :

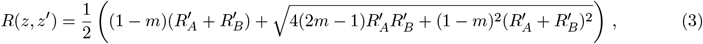

with

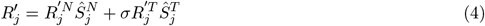

the mutant reproduction number in the population *j* (*A* or *B*), where

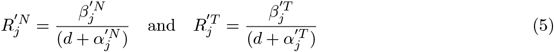

are respectively the mutant reproduction numbers in naive or vaccinated hosts in population *j*. In other words, the reproduction numbers 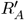 and 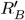 quantify the overall productivity of the mutant in each population. Note that the invasion fitness of the mutant, *R*(*z, z*^*′*^), depends both on the resident trait through susceptible densities at equilibrium 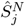and 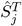 (included in 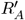 and 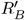), and on the mutant trait through the transmissibility (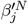 and 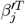) and virulence (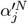 and 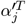) traits, which are included in 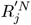 and 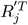 . The mutant invades if *R*(*z, z*^*′*^) *>* 1, and dies out if *R*(*z, z*^*′*^) ≤ 1. For completeness, we note that expression (3) is also valid for a mutant arising in a polymorphic resident population (i.e. when the trait *z* is a vector), as the resident environment only acts through the densities of susceptible hosts.

### 3.2 Selection gradient

In order to gain analytical insight on the long-term evolution of the trait of interest, *z* which governs the virulence and the transmission of the pathogen, we calculate the selection gradient, 𝒮. This amounts to calculating the partial derivative of the mutants invasion fitness *R*(*z, z*^*′*^) with respect to *z*^*′*^ evaluated at *z* = *z*^*′*^. The zeros of the selection gradient correspond to the evolutionary singularities, noted *z*^∗^. In our model, it can be expressed as (*Supplementary Information S*.*2*) :

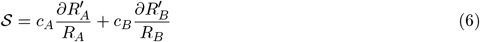

where 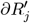 is a short-hand notation for the derivative of the mutant reproduction number in population *j*, calculated with respect to *z*^*′*^ and evaluated at *z*^*′*^ = *z* (see *Supplementary Information S*.*2* for a full expression), *R*_*A*_ and *R*_*B*_ are the local reproduction numbers in the resident population at equilibrium (defined as in equation (4)), and

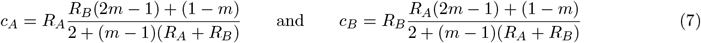

are the class reproductive values of pathogens in populations *A* and *B* respectively. The class reproductive values sum to 1 (e.g. *c*_*A*_ + *c*_*B*_ = 1, see *Supplementary Information S*.*2*) and can therefore be interpreted as measures of host quality for the pathogen which weigh the effects of selection in the two populations (Rousset, 1999; Gandon, 2004; Lion, 2018; Walter and Lion, 2021; Lion and Gandon, 2021). When the two populations are isolated (*m* = 0), there are two distinct singularities for each population, 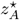 and 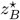, which corresponds to the zeros of 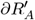 and 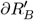 respectively. In the extreme case where a population (say, *A*) represents a habitat of very poor quality for pathogens (e.g. *c*_*A*_ tends towards 0), then, at the metapopulation level, the singularity *z*^⋆^ will tend towards the singularity of the good habitat, 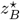 . On the other hand, when the two populations have similar qualities, the singularity at the metapopulation level will be intermediate between 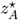 and 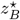 .

Using equation (7), we can rewrite condition 𝒮 = 0 as

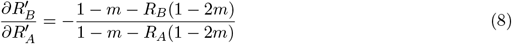

Equation (8) is reminiscent of *fitness set* representation that have been used to obtain a geometric understanding of adaptation in heterogeneous environments (Levins, 1962; Gandon et al., 2003), although here the graphical analysis is complicated by the fact that the pathogen’s fitness set is not fixed because it depends on epidemiological feedbacks via the densities of susceptible hosts. Yet, this expression can be used to understand how the evolutionary outcome depends on the interplay between the local productivity *R*_*j*_ of each population and migration. When migration is maximal (*m* = 1*/*2), the right-hand side of (8) is equal to −1, so that, at an evolutionary singularity, an increase in pathogen reproductive success in population *B* must be exactly compensated by a decrease in population *A*. However, when *m* decreases, the balance will depend on migration and on the relative productivity of the two populations. If a population has a higher productivity (i.e. *R*_*B*_ *> R*_*A*_) it will drive evolution towards a specialization to this population (i.e. *z*^⋆^ evolves towards 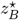). In other words, this *source-sink* evolutionary dynamics is driven by the source. Yet, since 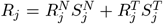, epidemiological feedbacks can affect the number of singularities and their positions with respect to 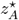 and 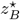, as we shall see in our numerical analysis below.

### 3.3 Stability of evolutionary equilibria

After convergence by successive mutations towards an evolutionary singularity, two different outcomes are possible. If the singularity cannot be invaded by nearby mutants, it corresponds to an evolutionarily stable strategy (ESS). If nearby mutants can invade, the singularity is called a branching point (BP), and the population can split into two coexisting morphs. The stability of the singularity can be investigated by calculating the second-order derivative of the invasion fitness, 𝒟, near the singularity *z*^⋆^. If 𝒟 *<* 0 the evolutionary singularity is stable. If 𝒟 *>* 0, the singularity is unstable. In our model, 𝒟 takes the form

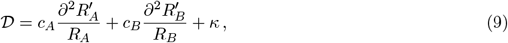

where 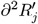 is the second order derivative of the mutant reproduction number in population *j* with respect to *z*^*′*^ evaluated at *z* = *z*^*′*^ (see *Supplementary Information S*.*2* for a full expression), and

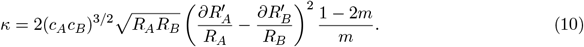

It is useful to interpret equations (9) by using the analogy of the fitness landscape. The first two terms on the right-hand side of equation (9) are a weighted average of the local curvature of the fitness landscapes in populations *A* and *B*, while *κ* in equation (10) depends on the squared difference of the slopes of the fitness landscapes, locally near the singularity. Thus, *κ* is always positive, and therefore acts as a destabilising force. In particular, this means that, even if selection is locally stabilising around the singularity *z*^⋆^ in both populations, so that the partial derivatives in equation (9) are both negative, selection can be disruptive at the metapopulation level when *κ* is large enough. This effect is due to divergent directional selection on the trait in the two populations, as captured by the difference 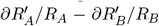. In general, the singular strategies will be evolutionarily stable if 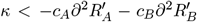 and unstable otherwise.

With a concave-down, monotonically increasing transmission-virulence trade-off, the sign of the second derivatives 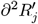 is negative unless the vaccine has a strong anti-growth component *r*_2_ (Gandon et al., 2003). Hence, when *κ* tends towards zero, we expect the evolutionarily singularity *z*^⋆^ to be always an ESS. From equation (10), we see that *κ* will be vanishingly small when (1) the class reproductive value of the pathogen is zero in one of the populations (e.g. *c*_*A*_*c*_*B*_ = 0); (2) the direction of selection is the same in both habitats (e.g. ∂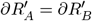); and (3) the metapopulation is well-mixed (*m* = 1*/*2).

On the other hand, equation (10) shows that a low migration rate will cause *κ* to increase and lead to evolutionarily unstable singularities. Hence, the singularity will tend to be unstable at low migration rates, unless the two populations resemble a single population (either because directional selection is equal in both 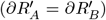 or because one of the populations does not contribute to selection (*c*_*A*_ = 0 or *c*_*B*_ = 0)).

### 3.4 Numerical exploration of the effect of migration and spatial heterogeneity

So far, our general adaptive dynamics analysis of pathogen evolution in a monomorphic resident population predicts that, in general, we expect a high migration rate to lead to a single ESS, whereas unstable singularities, potentially leading to polymorphism, can exist in the limit of low migration. In this section, we investigate different biological scenarios to make more specific predictions on the evolutionary outcomes, using both our monomorphic AD anaytical results and numerical simulations which allow us to explore the polymorphic dynamics of the population, and to relax the assumption of mutation-limited evolution used in AD.

We quantify the spatial heterogeneity of the metapopulation using the parameter *δ* = *ν*_*B*_ − *ν*_*A*_, which captures the variation in vaccination coverage between the two populations. Without loss of generality, we assume that the vaccination coverage is always higher in population *B*, so that *δ* takes values between 0 (homogeneous distribution, *ν*_*A*_ = *ν*_*B*_) and 1 (maximal heterogeneity, *ν*_*A*_ = 0 and *ν*_*B*_ = 1). For simplicity, we also assume in the main text that the average probability of vaccination at birth is *ν* = (*ν*_*A*_ + *ν*_*B*_)*/*2 = 0.5, so that *ν*_*A*_ ∈ [0, 0.5] and *ν*_*B*_ ∈ [0.5, 1]), an assumption that we relax in *Supplementary Information* S.7.

In line with previous theoretical studies on virulence evolution (Alizon et al., 2009), our results rely on a positive correlation between transmission and virulence, captured by an increasing and saturating transmission-virulence trade-off function. More specifically we use the functional form *β*(*z*) = *β*_0_*z/*(1 +*z*) in all our simulations.

#### Anti-infection and anti-transmission vaccines

For anti-infection (*r*_1_) or anti-transmission (*r*_3_) vaccines, we show that spatial structure does not affect pathogen evolution (*Supplementary Information S*.*2*). Indeed, even though the *r*_1_ and *r*_3_ components of the vaccine affect the efficacy of vaccination and the reproductive value of the pathogen in vaccinated hosts, the optimal virulence is independent of these parameters in the absence of coinfections (Gandon et al., 2001, 2003; Walter and Lion, 2021). Indeed, as discussed in previous studies in well-mixed single-population models the ES virulence is only affected by vaccination if the vaccine acts on virulence directly. Since the optimal virulence (i.e. the ES virulence in the absence of migration) does not vary between the two populations, the ES virulence depends neither on the migration rate *m* nor on the variation in vaccination coverage *δ*. In the following, we therefore focus on anti-growth (*r*_2_) and anti-virulence (*r*_4_) vaccines.

#### Anti-growth and anti-virulence vaccines

Non-spatial models have shown that anti-growth (*r*_2_) and anti-virulence (*r*_4_) vaccines can select for increased virulence (Gandon et al., 2001, 2003; Read et al., 2015; Walter and Lion, 2021). Our results show that, in line with these earlier results, both *r*_2_ and *r*_4_ vaccines select for higher virulence, but that this effect is modulated by the migration rate *m* and the heterogeneity in vaccination coverage *δ*. In particular, there is a marked difference between *r*_2_ and *r*_4_ vaccines: anti-growth vaccines (*r*_2_) allow for the coexistence of a lowand a high-virulence pathogen when migration is below a threshold (figures 2a and 3a). In contrast, anti-virulence vaccines (*r*_4_) are less conducive to the emergence of polymorphism (figures 2b and 3b).

**Figure 2:**
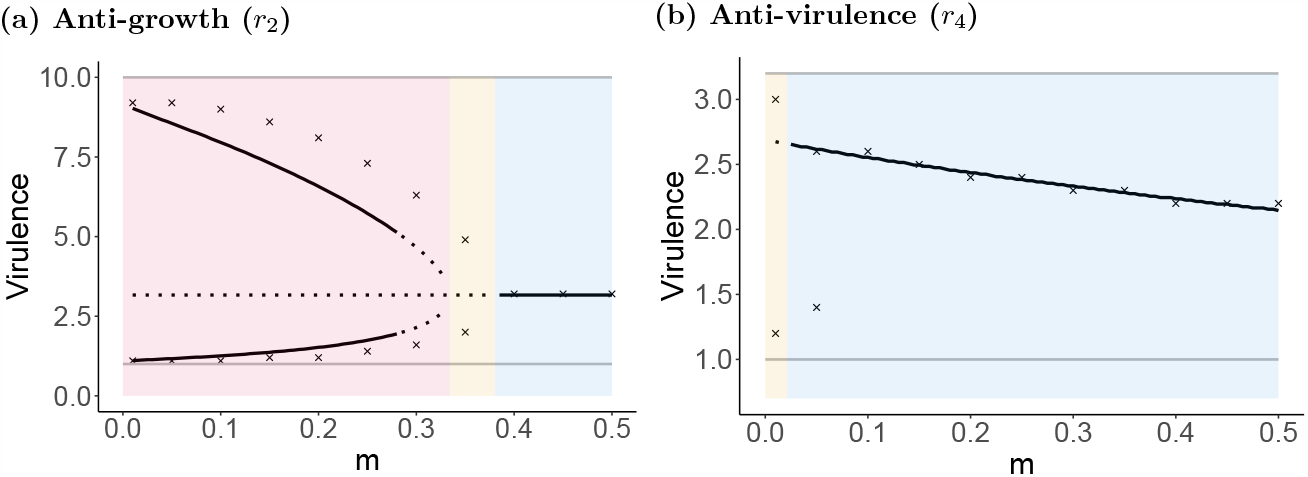
Effect of migration and vaccination on the long-term evolution of the pathogen. The plain and dotted black line represent the monomorphic singularities calculated by the adaptive dynamics. Plain black lines represents locally evolutionarily stable singularities while dotted black lines represent locally evolutionarily unstable singularities (branching points or repellor). The upper plain grey line represent the ESS in a population fully vaccinated, and the lower plain grey line represent the ESS in a population without vaccination. The black crosses represent the virulence strategies of the globally stable polymorphic equilibrium (see *Supplementary Information S*.*4* for further information). (a) For the anti-growth vaccine, when *m* ∈ {0, 0.33} we observe bistatility (pink area), with *m* ∈ {0.33, 0.38}, there is a unique branching point (yellow area), and with *m* ∈ {0.38, 0.5}, there is a unique ESS (blue area). For the anti-vrulence vaccine, when *m* ∈ {0, 0.02}, the monomorphic singularity is a branching point (yellow area) and with *m* ∈ {0.02, 0.5}, the singularity is an ESS (blue area). Here *δ* = 0.75, *μ* = 0.01, *V*_*m*_ = 0.001 and for (a) *r*_2_ = 0.9 and (b) *r*_4_ = 0.9 and all the other parameters as in **figure 1**.

For anti-growth vaccines (*r*_2_), our results predict that, at low migration, polymorphism arises due to local adaptation within the low-coverage and high-coverage habitats. As migration increases, the breakdown of spatial structure interferes with local adaptation, which favours a single generalist strategy. Overall, this corresponds to our general intuitive understanding of the interplay between migration and local adaptation in metapopulation models. More precisely, our AD analysis allows us to tease apart the effect of migration (*m*) vs heterogeneity (*δ*), and predicts three distinct long-term evolutionary ouctomes: (i) convergence towards a single ESS, (ii) evolutionary branching or (iii) evolutionary bistability, with convergence towards different ESSs depending on the initial conditions. Both outcomes (ii) and (iii) can potentially lead to polymorphism. To see this, we first explore the effect of the migration *m*, on the singularities, for a fixed value of the vaccination heterogeneity *δ* (figure 2). At high migration (*m* = 0.5), we have a single evolutionary stable virulence. As migration decreases and the two populations become less connected, the monomorphic singularity turns into a branching point, leading to the splitting of the population in two morphs that evolve towards a dimorphic ESS (the crosses in figure 2). As migration further decreases, we enter the bistability region, where the monomorphic singularity becomes an evolutionary repellor (central dotted line), surrounded by two other singularities. Near *m* = 0.3, these two singularities are branching points and the dynamics eventually converge towards the dimorphic ESS (crosses). For lower migration, the two singularities are evolutionarily stable, and evolutionary dynamics can evolve towards three distinct ESSs depending on the initial conditions: either one of two monomorphic ESSs (plain lines), or a dimorphic ESS (crosses). A similar pattern was observed in previous studies (Débarre et al., 2013; Mirrahimi and Gandon, 2020) that analysed the joint evolution of local adaptation and demography in a two-population model. Importantly, as in these previous studies, we find that the dimorphic ESS is always globally stable in the bistability region, whereas the monomorphic ESS are only locally stable (*Supplementary Information S*.*2*). Hence, with enough mutation and standing variation, the population dynamics will eventually converge to the globally stable dimorphic attractor. To numerically calculate the dimorphic ESS, we do not use the AD analysis, but rather directly use numerical simulations where we track the dynamics of a continuous trait distribution under selection and mutation (*Supplementary Infrormation* S.4). However, even if, for simplicity, we do not explicitly explore the dimorphic adaptive dynamics in our model, the AD analysis presented in the previous section is sufficient to shed light on the key factors (e.g. migration and habitat heterogeneity) that cause the shift from a single stable ESS to a dimorphic evolutionary outcome.

Interestingly, for anti-virulence (*r*_4_) vaccines, decreasing migration does not promote polymorphism, except when pathogen migration is completely local (figure 2b). We only observe a single ES virulence, which increases as migration becomes more local. This is because, in population B, which has a higher coverage of anti-virulence vaccine, infected hosts have a longer infection period, but no reduction in transmission. This leads to a higher productivity (*R*_*B*_ *> R*_*A*_) and a higher prevalence (see figure 1b). As discussed above, equation (8) shows that when migration becomes local, the evolutionary outcome is increasingly driven by the *source*. This explains why the ESS gets closer to the optimum of population *B*, which corresponds to higher virulence (Gandon et al., 2001, 2003). As explained in the Discussion, the discrepancy between the predictions for *r*_2_ and *r*_4_ vaccines can be explained by the specific epidemiological feedbacks that take place in each scenario.

In figure 3, we explore the interaction between the effects of the heterogeneity in vaccination coverage *δ* and migration on pathogen evolution, for both anti-growth and anti-virulence vaccines. In each of these figures, the bottom right corner corresponds to the most heterogeneous metapopulation (low migration and maximal difference in vaccination coverage). In contrast, the upper-left corner represents a completely homogeneous metapopulation (*δ* = 0 and *m* = 0.5), and corresponds to the results of Gandon et al. (2001) (without superinfection) or Walter and Lion (2021) (without temporal fluctuations). As expected, when *δ* increases (i.e. as the populations becomes more different), polymorphism emerges more readily, although again there is a clear difference between anti-growth and anti-virulence vaccines. We note however that this difference may be partly dependent on the specific transmission-virulence trade-off shape we assume in this study.

**Figure 3:**
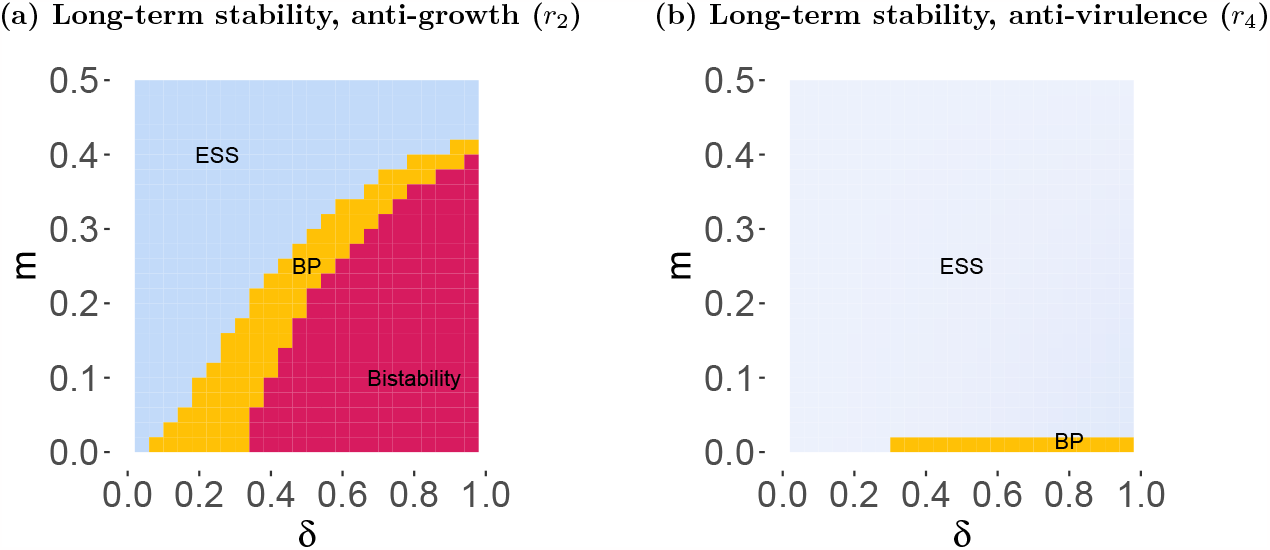
Effect of migration and heterogeneity in vaccination coverage on long-term evolutionary outcome. Long term evolutionary state calculated using adaptive dynamics, for (a) anti-growth and (b) anti-virulence vaccines. Evolutionary stable virulence (ESS) are represented in blue, following the gradient shown in **figure 4**, branching point (BP) in orange and bistability in pink, according to *m* and *δ*. Here *r*_*j*_ = 0.9 and the other parameters are as in **figure 1**.

In **figure 4** we explore the effects of a a vaccine that combines multiple components on the evolution of the pathogen. In particular, **figure 4a** shows that the combination of *r*_2_ and *r*_4_ effects can further expand the parameter space where polymorphism is expected to emerge and be stably maintained in the long-term. In **figure 4b** we also explore the effect of a vaccine with a combination of *r*_2_ and *r*_1_ effects. **Figure 4b** shows that even if *r*_1_ is expected to have no effect on pathogen evolution, we see a major interaction with the effect of *r*_2_ since *r*_1_ reduces the possibility of bistability and branching. A similar effect is observed in a combination between *r*_2_ and *r*_3_ (see *Supplementary Information S*.*3*).

**Figure 4:**
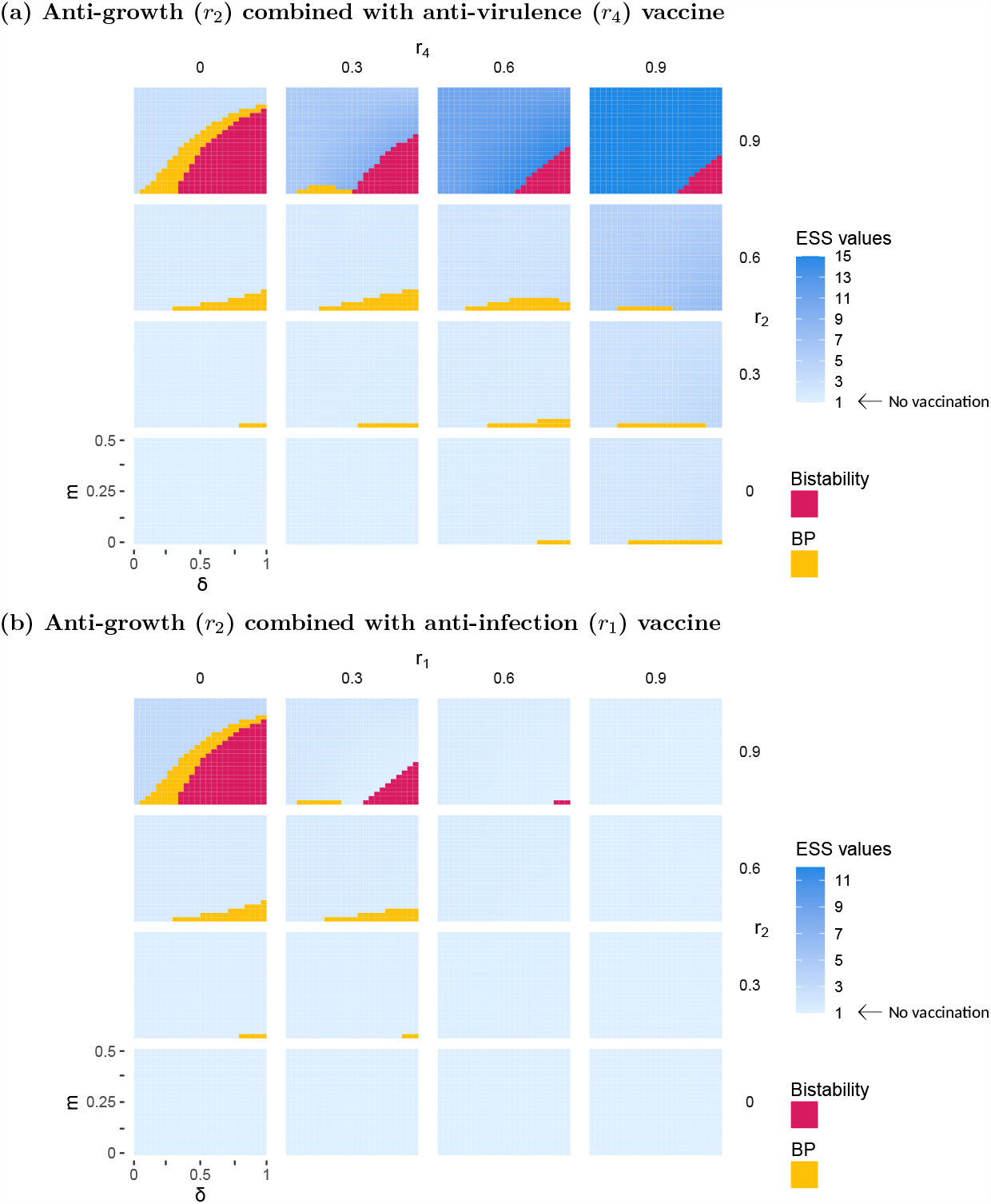
Long-term evolutionary outcome for different combination of vaccines. We plot the long-term evolutionary state calculated using adaptive dynamics, for anti-growth vaccine combined with anti-virulence, according to *m* and *δ*. Branching point (BP) are represented in yellow, bistability in pink, and ESS values in a gradient of blue. Other parameters as in **figure 1**.

## 4 Discussion

Here, we study the long-term evolution of pathogen life-history traits in a metapopulation, in response to the selective pressures caused by imperfect vaccines with different modes of action. Our analysis confirms the general message of earlier non-spatial models (Gandon et al., 2001, 2003; Walter and Lion, 2021) that there is a sharp dichotomy between vaccines that only have an effect on pathogen reproduction (e.g. anti-infection and anti-transmission vaccines) and vaccines that reduce the cost of virulence for the pathogen (i.e. anti-growth and anti-virulence vaccines). With the first type of vaccine, we predict no effect of vaccination on the long-term evolution of pathogen virulence (or potentially even a decrease in virulence when multiple infections are taken into account, see e.g. Gandon et al. (2001)). In contrast, for anti-growth and anti-virulence vaccines, we show that pathogen virulence can evolve in response to vaccination. In addition, we show that the long-term evolutionary outcome crucially depends on the degree of spatial heterogeneity of the metapopulation, which we quantify using two parameters: the migration rate *m* and the asymmetry in vaccine coverage *δ*. For high migration (*m* = 0.5) and low asymmetry (*δ* = 0), the metapopulation behaves like a single homogeneous population, and we recover the results of previous non-spatial models (Gandon et al., 2001, 2003; Walter and Lion, 2021). However, when migration is low and assymmetry is high, pathogen polymorphism becomes possible, leading to coexistence between two specialist strains, one that is adapted to the less vaccinated population, and the other that is adapted to the population with higher coverage. In other words, as in classical metapopulation models (Meszéna et al., 1997; Day, 2000; Ronce and Kirkpatrick, 2001; Débarre et al., 2013; Mirrahimi and Gandon, 2020), heterogeneity in selection (as measured by the asymmetry *δ*) and low migration favour the coexistence between genotypes locally adapted to each habitat. When the degree of heterogeneity of the metapopulation is below a threshold, our model predicts a single generalist ES strategy.

However, in contrast to classical local adaptation models which tend to assume Gaussian fitness functions maximized at fixed and externally determined optima, the effects we observe in our vaccination model are all mediated by epidemiological feedbacks, through the densities of vaccinated and unvaccinated infected hosts in each population. Interestingly, these epidemiological feedbacks act differently for anti-growth (*r*_2_) and anti-virulence (*r*_4_) vaccines: increasing the degree of heterogeneity in the metapopulation selects for more diversification with anti-growth vaccines, while polymorphism is only observed for maximal heterogeneity with anti-virulence vaccines. This occurs because *r*_4_ vaccines increase the duration of infection and boost the productivity of the highly vaccinated population which causes asymmetric migration between the two populations. In contrast, *r*_2_ vaccines do not increase the productivity of the highly vaccinated population because the increased duration of infections is balanced by the reduction in transmission. With *r*_4_ vaccines, the asymmetric influx of pathogens from the highly vaccinated population has two consequences on the long-term evolutionary outcome. First, it prevents the coexistence between two specialist genotypes because local selection to the unvaccinated population is swamped by migration from the vaccinated population. Second, asymmetric migration promotes adaptation to the more productive population (i.e. the highly vaccinated popualtion) and selects for high ES virulence. This source-sink interpretation is similar to the findings of earlier analyses of adaptation in two-population models (Meszéna et al., 1997; Day, 2000; Ronce and Kirkpatrick, 2001; Débarre et al., 2013; Mirrahimi and Gandon, 2020). Interestingly, this source-sink dynamics is very different with the anti-growth (*r*_2_) vaccine because pathogen prevalence drops in the vaccinated population. Hence, different vaccines have different consequences on the asymmetry between the populations. As discussed by Mirrahimi and Gandon (2020), this source-sink dynamics could be exacerbated by additional asymmetries in migration rates between populations but the exploration of these effects is beyond the scope of the present work.

It is interesting to compare the predictions of our two-patch model to previous analyses of the impact of spatial structure on the evolution of pathogen virulence. Using a lattice model, Zurita-Gutiérrez and Lion (2015) have shown that, for all types of vaccines, lower pathogen dispersal tends to select for lower ES virulence, except for high-efficacy anti-growth vaccines. In addition, they do not observe polymorphism. Although this prediction is consistent with the result that selection should favour more “prudent” pathogen strains in spatially structured populations (Haraguchi and Sasaki, 2000; van Baalen, 2002; Lion and van Baalen, 2008), it could appear at odds with our result that pathogen migration either has no or little effect on the ES virulence (for *r*_1_, *r*_2_ and *r*_3_ vaccines) or leads to increased ES virulence (for *r*_4_ vaccines) or to polymorphism. However, it is important to note that, although the model of Zurita-Gutiérrez and Lion (2015) does incorporate spatial structure by allowing a mixture of global and local transmission, it does not allow for heterogeneity in vaccination coverage as all new hosts have the same probability of being vaccinated, irrespective of their location. Furthermore, the effects observed in Zurita-Gutiérrez and Lion (2015) are mostly driven by kin competition for susceptible hosts. In our model, local populations are well-mixed and there is no kin selection.

Although the results in the main text assume a high vaccine efficacy (90%) and that the mean vaccination coverage at birth is 1/2 (and thus, (*ν*_*B*_ + *ν*_*A*_)*/*2 = 1*/*2), we have relaxed these assumptions in *Supplementary Information S*.*5*, where we generalise our results to all the possible combinations of vaccine coverage and efficacy. We show that an increase in pathogen migration, a decrease in vaccination asymmetry and a decrease in vaccine efficacy makes polymorphism less likely. This is consistent with our general message that, as the metapopulation becomes increasingly homogeneous, the potential for diversification weakens. In *Supplementary Information S*.*7*, we also show that a model where adults (and not newborns) are vaccinated yields qualitatively similar predictions.

There are several interesting potential extensions of this work. First, our model assumes that hosts either cannot recover, and therefore does not take into account natural immunity. We do not expect our qualitative results to be affected if recovery is total, but taking into account more realistic patterns of natural immunity would be technically more challenging and potentially impact our predictions. We note however that our model is a good approximation of many chronic diseases, and particularly of Marek’s disease in poultry, where evolution in response to vaccination has been well studied (Gimeno, 2008; Read et al., 2015). Second, our model does not account for coinfections and we know that within-host could potentially alter the effect of vaccination on the evolution of virulence (Gandon et al., 2001). Third, although we have framed our results in the context of vaccination against a pathogen infecting a single species of hosts, our analytical results are more general because they allow the pathogen’s life-history traits to depend on different types of host (*N* and *T* in the present model). This implies that the same analytical expressions capture the direction of selection and the condition of evolutionarily stability for a pathogen infecting two host species (Gandon, 2004). Moreover, the focus on vaccines is by no means exclusive, as the four modes of action we distinguish can also be used to describe the action of other forms of heterogeneity, such as genetic resistance in the host or other treatments. In the context of chronic diseases, there are potentially interesting parallels to draw with imperfect antiretroviral therapies in HIV, although the complexity of HIV dynamics would require an adaptation of our epidemiological assumptions. Third, a limitation of our study is that it relies on a separation of time scales assumption to make prediction on the long-term evolutionary outcome. However, for many practical purposes, what matters most is the joint dynamics of strain diversity and epidemiological dynamics over shorter time scales (Gandon and Lion, 2022). Hence, it would be interesting to extend this model to take into account both spatial heterogeneity and epidemic dynamics on the transient spread of new pathogen variants. Fourth, our adaptive dynamics framework is not well suited to capture the inherently stochastic nature of the spread of new mutations (Restif and Grenfell, 2006, 2007; Day et al., 2022; Gandon et al., 2022). In a similar two-patch stochastic model, Gerrish et al. (2021) have shown that heterogeneity in vaccine distribution promotes the emergence of vaccine escape mutants. It would be interesting to explore how demographic stochasticity could alter the emergence and the spread of new pathogen variants.

From an applied point of view, the consequences of the evolution of pathogen polymorphism on longterm infectious disease management are not obvious. On the one hand, our model does not predict that polymorphism affects the cumulative number of deaths (see *Supplementary Information S*.*6*), although this specific prediction is probably sensitive to epidemiological assumptions. On the other hand, the circulation of two variants instead of one is a major challenge for the design of effective control strategies. For instance, this may require the development of vaccines targeting all strains, or specifically the most virulent variants as in the vaccination against *Streptococcus pneumoniae* (Masomian et al., 2020; Musher et al., 2022) or against human papillomavirus (Schiller and Lowy, 2012; Koutsky and Harper, 2006). Further studies are thus required to investigate how strain coexistence, which is expected to be a qualitatively robust outcome of pathogen evolution in populations with a high degree of spatial heterogeneity, needs to be taken into account in public health strategies. In addition, our results suggest that the risk of high-virulence evolution can be mitigated by the use of vaccines that combine both anti-infection and anti-growth effects. This is reassuring as most vaccines are unlikely to squarely fall into one of the four categories we distinguish in this study.

At a conceptual level, there is a parallel between our present investigation of the effect of spatial heterogeneity in vaccination coverage, and our previous analysis of the effect of temporal heterogeneity through periodic vaccination coverage (Walter and Lion, 2021). In this previous study, the pathogen sequentially experiences two very different habitats (corresponding to maximal or minimal coverage respectively, which corresponds to *δ* = 1 in our spatial model, and the period of fluctuations allows for varying the amount of time spent in each habitat (much like the migration parameter *m*). Both studies predict that heterogeneity in vaccination coverage (whether it is spatial or temporal) has no effect on the ES virulence for anti-infection and anti-transmission vaccines. However, for anti-growth and antivirulence vaccines, the parallel between temporal and spatial heterogeneity is not straightforward. First, the model with periodic fluctuations in vaccination coverage never predicts strain coexistence. Second, for anti-virulence vaccine, increasing the period of temporal fluctuations leads to lower virulence, whereas decreasing migration leads to higher virulence in the spatial model. Hence, the effect of increased heterogeneity is markedly different in the spatial and temporal models. This calls for further exploration of the consequences of a combination of both spatial and temporal variations in vaccine coverage on the evolution of pathogen traits. For instance, in the context of the COVID-19 pandemic, Kortessis et al. (2020) have used a similar metapopulation model to study the effect of asynchronous public health measures on disease prevalence. A very interesting avenue for future research would be to extend this analysis to explore how spatio-temporal fluctuations of the environment affect the shortand long-term evolution of pathogens. Understanding both the epidemiological and the evolutionary consequences of different deployment strategies of vaccination may help identify more effective vaccination strategy that minimise both the cumulated number of cases and the speed at which the pathogen adapts to vaccination.

## Funding

This work was funded by grants ANR-16-CE35-0012 “STEEP” to SL and ANR-17-CE35-0012 “EVO-MALWILD” to SG from the Agence Nationale de la Recherche.

## Acknowledgements

We thank two anonymous reviewers for their critical reading and helpful comments.

## Codes

The code used to generate the figures (Mathematica and R scripts) can be found at https://zenodo.org/records/10212665.

## Notes

### Competing Interest Statement

The authors have declared no competing interest.

### Summary of Updates

- general clarification of the text - figures revised

