## Additional methods and figures for "Effect of unequal vaccination coverage and migration on long-term pathogen evolution in a metapopulation"

November 27, 2023

### S.1 Epidemiology

#### S.1.1 Full ODE system

With the description of the host-pathogen life cycle given in the main text, the metapopulation dynamics are described by the following 8-equation system:

$$\begin{aligned}
\frac{\mathrm{d}S_A^N}{\mathrm{d}t} &= b(1-\nu_A) - S_A^N(d+H_A)\,,\\
\frac{\mathrm{d}S_A^T}{\mathrm{d}t} &= \nu_A b - S_A^T(d+\sigma H_A)\,,\\
\frac{\mathrm{d}S_B^N}{\mathrm{d}t} &= b(1-\nu_B) - S_B^N(d+H_B)\,,\\
\frac{\mathrm{d}S_B^T}{\mathrm{d}t} &= \nu_B b - S_B^T(d+\sigma H_B)\,,\\
\frac{\mathrm{d}I_A^N}{\mathrm{d}t} &= S_A^N H_A - (d+\alpha_A^N)I_A^N\,,\\
\frac{\mathrm{d}I_A^T}{\mathrm{d}t} &= \sigma S_A^T H_A - (d+\alpha_A^T)I_A^T\,,\\
\frac{\mathrm{d}I_B^N}{\mathrm{d}t} &= S_B^N H_B - (d+\alpha_B^N)I_B^N\,,\\
\frac{\mathrm{d}I_B^T}{\mathrm{d}t} &= \sigma S_B^T H_B - (d+\alpha_B^T)I_B^T\,,
\end{aligned} \tag{S.1}$$

where

$$H_A = (1-m)h_A + mh_B$$

is the force of infection experienced by susceptible hosts in population $A$ from either local infected hosts (with probability $1-m$) or infected hosts of the other population (with probabilty $B$). Similarly, we have

$$H_B = (1-m)h_B + mh_A$$

#### S.1.2 Dynamics

A detailed analysis of the epidemiological dynamics is beyond the scope of this article, but typical dynamics for an anti-growth or an anti-virulence vaccine are given in **figure S.1**.

**(a) Anti-growth**

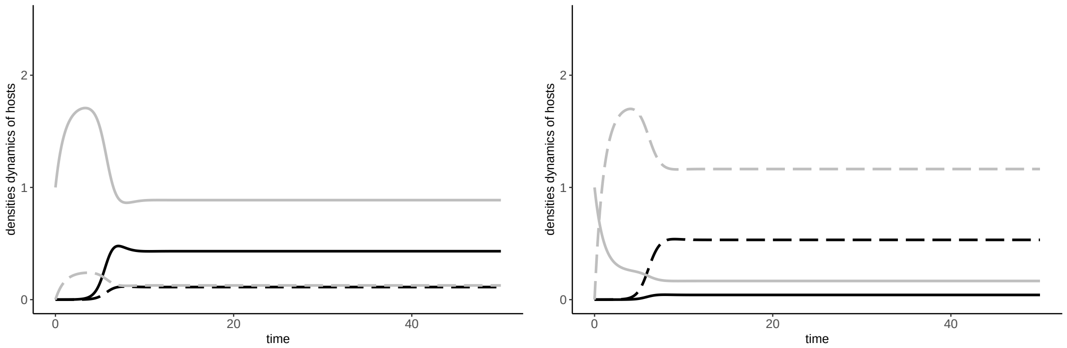


**(b) Anti-virulence**

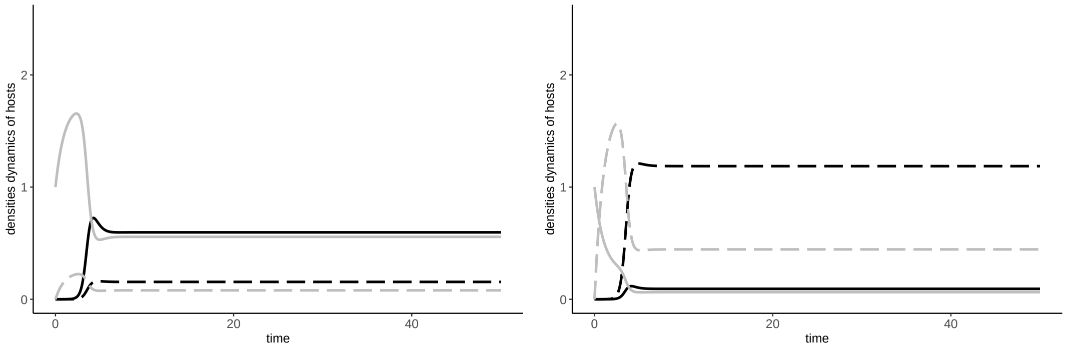


**Figure S.1:** Dynamics of the model with (a) anti-growth and (b) anti-virulence vaccines, in the population $A$ (left panels) or $B$ (right panels). We have represented the naive susceptible in plain grey, the vaccinated susceptible in dashed grey, the naive infected in plain black and the vaccinated infected in dashed black. Here, $b = 2$, $d = 1$, $z = 1$, $\delta = 0.75$, $m = 0.2$ and $r_j = 0.9$.

#### S.1.3 Equilibrium

In a disease-free environment, the densities of susceptible hosts settle at an equilibrium ($S_{0A}^N$, $S_{0A}^T$, $S_{0B}^N$, $S_{0B}^T$), that can be calculated as :

$$\left(\frac{(1-\nu_A)b}{d}, \frac{\nu_A b}{d}, \frac{(1-\nu_B)b}{d}, \frac{\nu_B b}{d}\right) \tag{S.2}$$

However, we do not have explicit expression for an the endemic equilibrium.

#### S.1.4 Basic reproduction number

We use the Next Generation theorem to calculate the pathogen basic reproduction number in the metapopulation (Diekmann et al., 1990; Hurford et al., 2009; Lion and Metz, 2018). From the dynamics of infected hosts, we have

$$\frac{\mathrm{d}}{\mathrm{d}t}\begin{pmatrix} I_A^N \\ I_A^T \\ I_B^N \\ I_B^T \end{pmatrix} = (\mathbf{F} - \mathbf{V})\begin{pmatrix} I_A^N \\ I_A^T \\ I_B^N \\ I_B^T \end{pmatrix}$$

where

$$\mathbf{F} = \begin{pmatrix} (1-m)S_{0A}^N\beta_A^N & (1-m)S_{0A}^N\beta_A^T & mS_{0A}^N\beta_B^N & mS_{0A}^N\beta_B^T \\ \sigma(1-m)S_{0A}^T\beta_A^N & \sigma(1-m)S_{0A}^T\beta_A^T & \sigma mS_{0A}^T\beta_B^N & \sigma mS_{0A}^T\beta_B^T \\ mS_{0B}^N\beta_A^N & mS_{0B}^N\beta_A^T & (1-m)S_{0B}^N\beta_B^N & (1-m)S_{0B}^N\beta_B^T \\ \sigma mS_{0B}^T\beta_A^N & \sigma mS_{0B}^T\beta_A^T & \sigma(1-m)S_{0B}^T\beta_B^N & \sigma(1-m)S_{0B}^T\beta_B^T \end{pmatrix}$$

is the transmission matrix and

$$\mathbf{V} = \begin{pmatrix} -(d+\alpha_A^N) & 0 & 0 & 0 \\ 0 & -(d+\alpha_A^T) & 0 & 0 \\ 0 & 0 & -(d+\alpha_B^N) & 0 \\ 0 & 0 & 0 & -(d+\alpha_B^T) \end{pmatrix}$$

is the mortality matrix. It is straightforward to check that the conditions of the NGT are satisfied (Hurford et al., 2009), so that the basic reproduction number is given by the dominant eigenvalue of matrix $\mathbf{F}.\mathbf{V}^{-1}$ (Diekmann et al., 1990; Hurford et al., 2009)

$$\mathbf{F}.\mathbf{V}^{-1} = \begin{pmatrix} (1-m)S_{0A}^N R_A^N & (1-m)S_{0A}^N R_A^T & mS_{0A}^N R_B^N & m\beta_B^T S_{0A}^N R_B^T \\ \sigma(1-m)S_{0A}^T R_A^N & \sigma(1-m)S_{0A}^T R_A^T & \sigma m S_{0A}^T R_B^N & \sigma m S_{0A}^T R_B^T \\ mS_{0B}^N R_A^N & mS_{0B}^N R_A^T & (1-m)S_{0B}^N R_B^N & (1-m)S_{0B}^N R_B^T \\ \sigma m S_{0B}^T R_A^N & \sigma m S_{0B}^T R_A^T & \sigma(1-m)S_{0B}^T R_B^N & \sigma(1-m)S_{0B}^T R_B^T \end{pmatrix}. \quad \text{(S.3)}$$

where $R_k^N = \beta_k^N/(d+\alpha_k^N)$ and $R_k^T = \beta_k^T/(d+\alpha_k^T)$. Thus, we can calculate $R_0$ as

$$R_0 = \frac{1}{2}\left((1-m)(R_{0A}+R_{0B}) + \sqrt{4(2m-1)R_{0A}R_{0B} + (1-m)^2(R_{0A}+R_{0A})^2}\right). \quad \text{(S.4)}$$

with $R_{0k} = R_k^N S_{0k}^N + \sigma R_k^T S_{0k}^T$, the basic reproduction number in population $k$, and the disease-free equilibria $S_{0k}^N$ and $S_{0k}^T$ are given in equation S.2.

### S.2 Evolution

#### S.2.1 Mutant invasion fitness

A proxy for the mutant invasion fitness can be calculated following the method used to calculate the basic reproduction ratio of the pathogen, except that we now calculate it in the resident endemic equilibrium (Lion and Metz, 2018). Hence, we calculate the dominant eigenvalue of the matrix $\mathbf{F}'.\mathbf{V}'^{-1}$, where $\mathbf{F}'$ and $\mathbf{V}'$ are analogous to $\mathbf{F}$ and $\mathbf{V}$ in section S.1.4 except that the disease-free equilibria of susceptible hosts are replaced by the corresponding endemic equilibria. We obtain:

$$R(z,z') = \frac{1}{2}\left((1-m)(R'_A+R'_B) + \sqrt{4(2m-1)R'_A R'_B + (1-m)^2(R'_A+R'_B)^2}\right), \quad \text{(S.5)}$$

where the definitions of $R'_A$ and $R'_B$ are given in the main text.

#### S.2.2 Selection gradient expression

The selection gradient corresponds to the derivative of the mutant invasion fitness $R(z,z')$ with respect to $z'$, evaluated at $z = z'$. We have

$$\mathcal{S} = \left.\frac{\partial R(z,z')}{\partial z'}\right|_{z'=z}.$$

For a neutral mutant, we have $R(z,z) = 1$ which leads to the following equality :

$$\sqrt{4(2m-1)(R_A R_B) + (1-m)^2(R_A+R_B)^2} = 2 - (1-m)(R_A+R_B) \quad \text{(S.6)}$$

which can be used to rewrite the selection gradient as

$$\mathcal{S} = \left(\frac{1-m}{2}\right)(\partial R'_A + \partial R'_B) + \frac{1}{4}\frac{4(2m-1)(\partial R'_A R_B + R_A \partial R'_B) + 2(1-m)^2(R_A+R_B)(\partial R'_A + \partial R'_B)}{2-(1-m)(R_A+R_B)}. \quad \text{(S.7)}$$

After rearrangement, we obtain

$$\mathcal{S} = X_A \partial R'_A + X_B \partial R'_B \quad \text{(S.8)}$$

with

$$X_A = \frac{(2m-1)R_B + (1-m)}{2-(1-m)(R_A+R_B)},$$
$$X_B = \frac{(2m-1)R_A + (1-m)}{2-(1-m)(R_A+R_B)}. \tag{S.9}$$
(S.10)

From eq.S.6, we can further derive the following equalities:

$$\begin{aligned}
\Leftrightarrow 4(2m-1)(R_AR_B) + (1-m)^2(R_A+R_B)^2 &= (2-(1-m)(R_A+R_B))^2 \\
\Leftrightarrow 4(2m-1)(R_AR_B) + (1-m)^2(R_A+R_B)^2 &= 4 + (1-m)^2(R_A+R_B)^2 - 4(1-m)(R_A+R_B) \\
\Leftrightarrow (1-m)(R_A+R_B) + (2m-1)R_AR_B = 1 &
\end{aligned} \tag{S.11}$$

The last equality can be used to show that

$$\begin{aligned}
X_AR_A + X_BR_B &= \frac{2(2m-1)R_AR_B + (1-m)(R_A+R_B)}{2-(1-m)(R_A+R_B)} \\
&= \frac{2\left(1-(1-m)(R_A+R_B)\right) + (1-m)(R_A+R_B)}{2-(1-m)(R_A+R_B)}, \\
&= 1
\end{aligned} \tag{S.12}$$

We can then define $c_A = X_AR_A$ and $c_B = X_BR_B$ the class reproductive values normalised such that $c_A + c_B = 1$ (Lion, 2018; Walter and Lion, 2021).

#### S.2.3 Second order derivative calculation

The second order derivative informs about the evolutionarily stability of singularities and is given by

$$\mathcal{D} = \left.\frac{\partial^2 R(z,z')}{\partial z'^2}\right|_{z'=z=z^*}. \tag{S.13}$$

If $\mathcal{D} < 0$ the evolutionary singularity is evolutionarily stable (ESS). If $\mathcal{D} > 0$, the singularity is unstable (branching point). In our model, we obtain :

$$\begin{aligned}
\mathcal{D} = \frac{1}{2}\Big(&(1-m)(\partial^2 R'_A + \partial^2 R'_B) \\
&+ \frac{4(2m-1)(\partial^2 R'_A R_B + \partial^2 R'_B R_A + 2\partial R'_A \partial R'_B) + 2(1-m)^2\left((\partial R'_A + \partial R'_B)^2 + (R_A+R_B)(\partial^2 R'_A + \partial^2 R'_B)\right)}{2\sqrt{4(2m-1)R_AR_B + (1-m)^2(R_A+R_B)^2}} \\
&- \frac{(4(2m-1)(\partial R'_A R_B + R_A \partial R'_B) + 2(1-m)^2(R_A+R_B)(\partial R'_A + \partial R'_B))^2}{4(4(2m-1)R_AR_B + (1-m)^2(R_A+R_B)^2)^{3/2}}\Big).
\end{aligned} \tag{S.14}$$

Expression (S.14) can be rewritten as $\mathcal{D} = X_A\partial^2 R'_A + X_B\partial^2 R'_B + \kappa$, with :

$$\kappa = \frac{2m^2(1-2m)(\partial R'_A - \partial R'_B)^2}{(2-(1-m)(R_A+R_B))^3}. \tag{S.15}$$

We have

$$\begin{aligned}
X_AX_B &= \frac{(1-m)^2 + (1-2m)^2R_AR_B - (1-2m)(1-m)(R_A+R_B)}{(2-(1-m)(R_A+R_B))^2}, \\
&= \frac{(1-m)^2 - (1-2m)}{(2-(1-m)(R_A+R_B))^2}, \\
&= \frac{m^2}{(2-(1-m)(R_A+R_B))^2}
\end{aligned} \tag{S.16}$$

We can then rewrite $\kappa$ as :

$$\kappa = 2(X_AX_B)^{3/2}\frac{1-2m}{m}\left(\partial R'_A R_B - \partial R'_B R_A\right)^2. \tag{S.17}$$

Using $c_A = X_A R_A = 1 - c_B = 1 - X_B R_B$, we can express the second order derivative as

$$\mathcal{D} = c_A \frac{\partial^2 R'_A}{R_A} + c_B \frac{\partial^2 R'_B}{R_B} + 2(c_A c_B)^{3/2}\sqrt{R_A R_B}\left(\frac{\partial R'_A}{R_A} - \frac{\partial R'_B}{R_B}\right)^2 \frac{1-2m}{m} \tag{S.18}$$

which correspond to the equation 9 in the main text.

In addition, it is useful to note that, when the two subpopulations are completely isolated, we have $R_A = 1$ and $R_B = 1$, so that, for small migration rates, we expect $R_A = 1 + O(m)$ and $R_B = 1 + O(m)$. Hence, plugging this into the expressions for $c_A$ and $c_B$, we have

$$c_A c_B = \frac{1}{4} + O(m).$$

#### S.2.4 $R_A$ and $R_B$ and their derivatives

In the above calculations, we use the notations $R'_A$ and $R'_B$ to characterise the mutant reproductive numbers in the population $A$ or $B$ as

$$\begin{aligned} R'_A &= \hat{S}^N_A R^N_A + \sigma \hat{S}^T_A R^T_A = \hat{S}^N_A \frac{\beta^N_A[z']}{d + \alpha^N_A[z']} + \sigma \hat{S}^T_A \frac{\beta^T_A[z']}{d + \alpha^T_A[z']}, \\ R'_B &= \hat{S}^N_B R^N_B + \sigma \hat{S}^T_B R^T_B = \hat{S}^N_B \frac{\beta^N_B[z']}{d + \alpha^N_B[z']} + \sigma \hat{S}^T_B \frac{\beta^T_B[z']}{d + \alpha^T_B[z']}. \end{aligned} \tag{S.19}$$

We use the notation $\partial R'_A$ and $\partial R'_B$ for the first order derivative of $R'_A$ and $R'_B$ with respect to the mutant strain $z'$, evaluated at $z = z'$ as

$$\begin{aligned} \partial R'_A &= \hat{S}^N_A \left(\frac{(d+\alpha^N_A)\partial\beta^N_A - \beta^N_A \partial\alpha^N_A}{(d+\alpha^N_A)^2}\right) + \sigma \hat{S}^T_A \left(\frac{(d+\alpha^T_A)\partial\beta^T_A - \beta^T_A \partial\alpha^T_A}{(d+\alpha^T_A)^2}\right), \\ \partial R'_B &= \hat{S}^N_B \left(\frac{(d+\alpha^N_B)\partial\beta^N_B - \beta^N_B \partial\alpha^N_B}{(d+\alpha^N_B)^2}\right) + \sigma \hat{S}^T_B \left(\frac{(d+\alpha^T_B)\partial\beta^T_B - \beta^T_B \partial\alpha^T_B}{(d+\alpha^T_B)^2}\right). \end{aligned} \tag{S.20}$$

We use the notation $\partial^2 R'_A$ and $\partial^2 R'_B$ for the second order derivative of $R'_A$ and $R'_B$, with respect to the mutant strain $z'$, evaluated at $z = z'$ as

$$\begin{aligned} \partial^2 R'_A &= \hat{S}^N_A \left(\frac{2\beta^N_A(\partial\alpha^N_A)^2 - (d+\alpha^N_A)\partial^2\alpha^N_A}{(d+\alpha^N_A)^3} - \frac{2\partial\alpha^N_A\partial\beta^N_A}{(d+\alpha^N_A)^2} + \frac{\partial^2\beta^N_A}{d+\alpha^N_A}\right) + \\ &\quad \sigma \hat{S}^T_A \left(-\frac{\beta^T_A((d+\alpha^T_A)\partial^2\alpha^T_A - 2(\partial\alpha^T_A)^2)}{(d+\alpha^T_A)^3} - \frac{2\partial\alpha^T_A\partial\beta^T_A}{(d+\alpha^T_A)^2} + \frac{\partial^2\beta^T_A}{d+\alpha^T_A}\right), \\ \partial^2 R'_B &= \hat{S}^N_B \left(\frac{2\beta^N_B(\partial\alpha^N_B)^2 - (d+\alpha^N_B)\partial^2\alpha^N_B}{(d+\alpha^N_B)^3} - \frac{2\partial\alpha^N_B\partial\beta^N_B}{(d+\alpha^N_B)^2} + \frac{\partial^2\beta^N_B}{d+\alpha^N_B}\right) + \\ &\quad \sigma \hat{S}^T_B \left(-\frac{\beta^T_B((d+\alpha^T_B)\partial^2\alpha^T_B - 2(\partial\alpha^T_B)^2)}{(d+\alpha^T_B)^3} - \frac{2\partial\alpha^T_B\partial\beta^T_B}{(d+\alpha^T_B)^2} + \frac{\partial^2\beta^T_B}{d+\alpha^T_B}\right), \end{aligned} \tag{S.21}$$

#### S.2.5 Anti-infection and anti-transmission vaccines

Anti-infection vaccines aim to reduce host susceptibility to pathogens and anti-transmission vaccines reduce the transmission of infected to susceptible hosts. With these vaccines, we have the following measure of pathogen fitness in population $A$ :

$$R'_A = \frac{\beta'^N_A}{d + \alpha'^N_A}\left(\hat{S}^N_A + (1-r_1)(1-r_3)\hat{S}^T_A\right).$$

Setting $Z_A = \hat{S}_A^N + (1-r_1)(1-r_3)\hat{S}_A^T$, it simplifies to $R'_A = R_A'^N Z_A$.

The only difference between subpopulations $A$ and $B$ is their vaccine coverage, and therefore naive hosts have the same transmission and mortality. Thus $R_A'^N = R_B'^N = R'^N$ and the selection gradient eq. 6 in the main text can be expressed as :

$$\mathcal{S} = \frac{\partial R'^N}{R^N}(Z_A c_A + Z_B c_B) \tag{S.22}$$

Because the susceptible densities ($Z_k$) and class reproductive values ($c_k$) are positive, $\mathcal{S} = 0$ is equivalent to $\partial R'^N = 0$. Finding the evolutionarily singularities $z^*$ therefore boils down to maximizing $\beta'^N/(d+\alpha'^N)$. Hence, $z^*$ is independent of $r_1$ and $r_3$, and therefore anti-infection and anti-transmission vaccines do not affect long-term the evolutionary outcome.

#### S.2.6 Pairwise Invasibility Plots and density plots

Under adaptive dynamics assumptions, starting from a monomorphic resident population, there are three different types of monomorphic evolutionary singularities in our model of pathogen evolution. First, we can observe an evolutionary stable virulence (ESS, **figure S.2**), that is, when a mutant reaches this singularity, it cannot be invaded by another mutant strain. Second, the singularity can be a branching point (BP, **figure S.3**, that is it can be invaded by neighbouring mutants, and potentially lead to polymorphism. Third, we can observe a bistability (**figure S.3**), that is, a repellor surrounded by two ESSs. Note however that this analysis is local: while the two ESSs are locally stable in the bistable case, there is also a global stable polymorphic equilibrium (Débarre et al., 2013; Mirrahimi and Gandon, 2020). Hence, numerical simulations with continuous trait distributions will typically converge towards a dimorphic equilibrium distribution instead of converging towards a unique ESS depending on the initial condition (as predicted by AD).

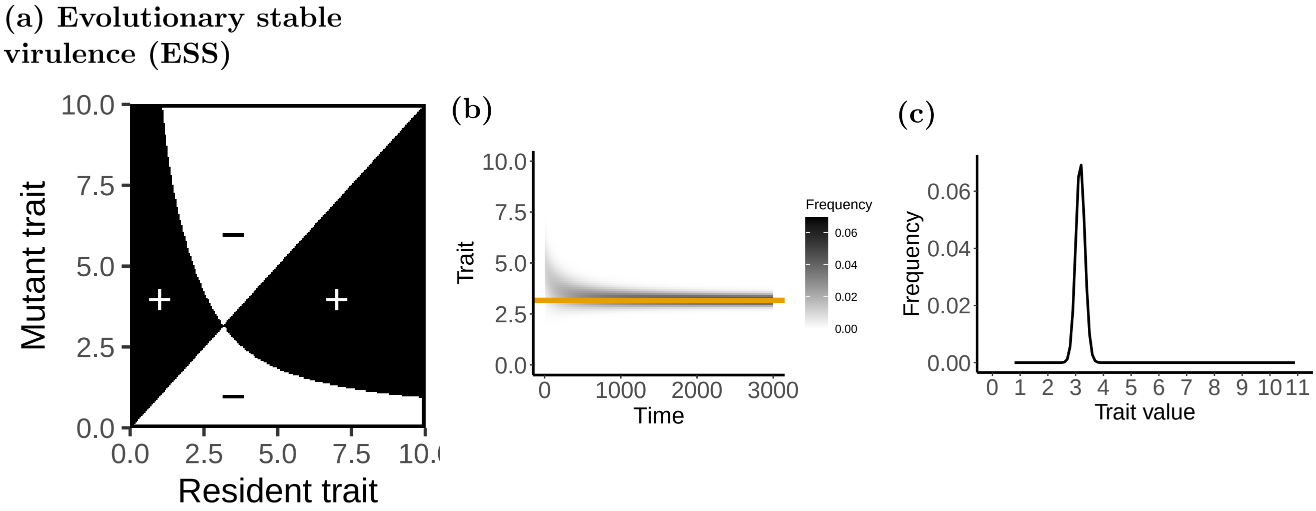


**Figure S.2:** Evolutionary stable virulence (ESS) for an anti-growth vaccine. This corresponds to $m = 0.45$ and $\delta = 0.75$ in **figure 3** in the main text. (a) Pairwise Invasibility Plot where the mutant strain wins the competition in white areas and the resident strain wins in black areas. (b) Simulated evolutionary tree in a polymorphic population (see Appendix S.4). The orange dotted line shows the ESS predicted by adaptive dynamics. (c) Virulence distribution in a polymorphic population. In (b) and (c), the initial distribution of trait centred on $z = 5$ and mutations are rare and have small effect ($\mu = 0.001$, $V_m = 0.01$). Here $r_2 = 0.9$, and other parameters are as in **figure 1** in the main text.

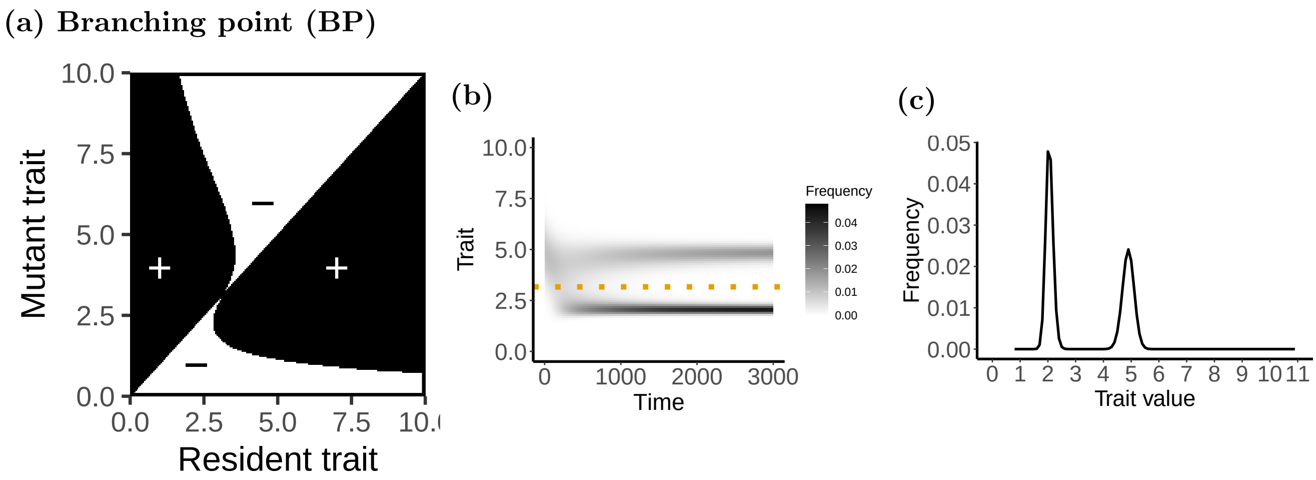


**Figure S.3:** Branching point (BP) for an anti-growth vaccine. This corresponds to $m = 0.35$ and $\delta = 0.75$ in **figure 3** in the main text. (a) Pairwise Invasibility Plot where the mutant strain wins the competition in white areas and the resident strain wins in black areas. (b) Simulated evolutionary tree in a polymorphic population (see Appendix S.4). The orange dotted line shows the branching point predicted by adaptive dynamics. (c) Virulence distribution in a polymorphic population. In (b) and (c), the initial distribution of trait centred on $z = 5$ and mutations are rare and have small effects ($\mu = 0.001$, $V_m = 0.01$). Here $r_2 = 0.9$, and other parameters are as in **figure 1** in the main text.

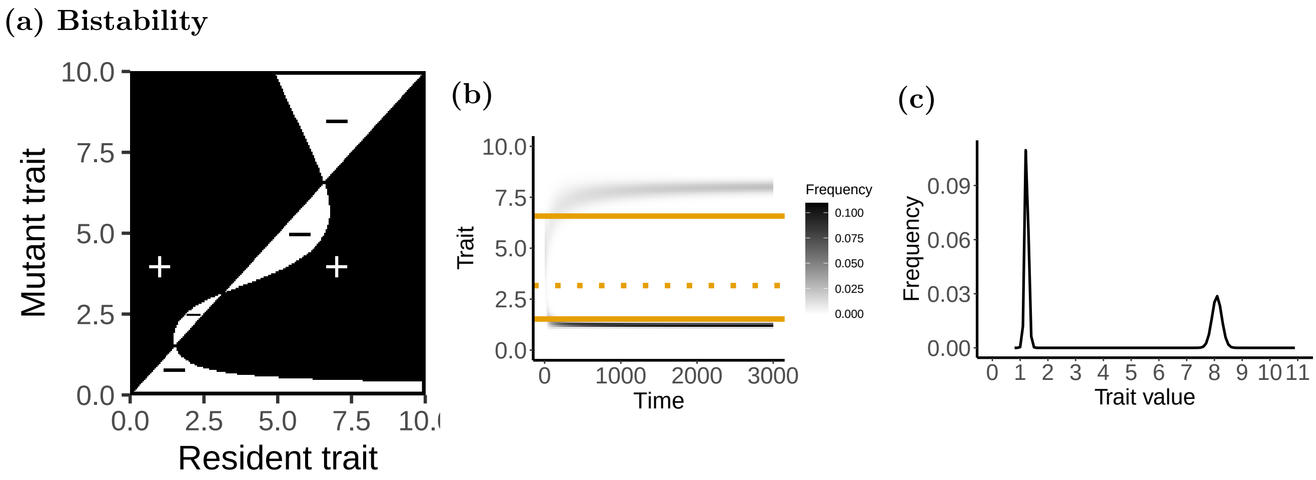


**Figure S.4:** Bistability for an anti-growth vaccine. This corresponds to $m = 0.2$ and $\delta = 0.75$ in **figure 3** in the main text. (a)Pairwise Invasibility Plot where the mutant strain wins the competition in white areas and the resident strain wins in black areas. (b) Simulated evolutionary tree in a polymorphic population (see Appendix S.4). The orange solid lines show the predicted monomorphic ESSs, the orange dotted line shows the predicted repellor, all calculated by adaptive dynamics. Note that the polymorphic simulatiosn converge towards the globally stable dimorphic ESS, and not towards one of the locally stable monomorphic ESSs. (c) Virulence distribution in a polymorphic population. In (b) and (c), the initial distribution of trait centred on $z = 5$ and the mutations are rare and have small effects ($\mu = 0.001$, $V_m = 0.01$). Here $r_2 = 0.9$, and other parameters are as in **figure 1** in the main text.

### S.3 Combination of vaccines

We have explored a combination of vaccine of different mode of action, $r_2$ with $r_1$ or $r_3$ (**figure S.5**), $r_4$ with $r_1$ or $r_3$ (**figure S.6**) and $r_2$ with $r_4$ (**figure S.7**).

**(a) Anti-growth ($r_2$) with anti-infection ($r_1$)**

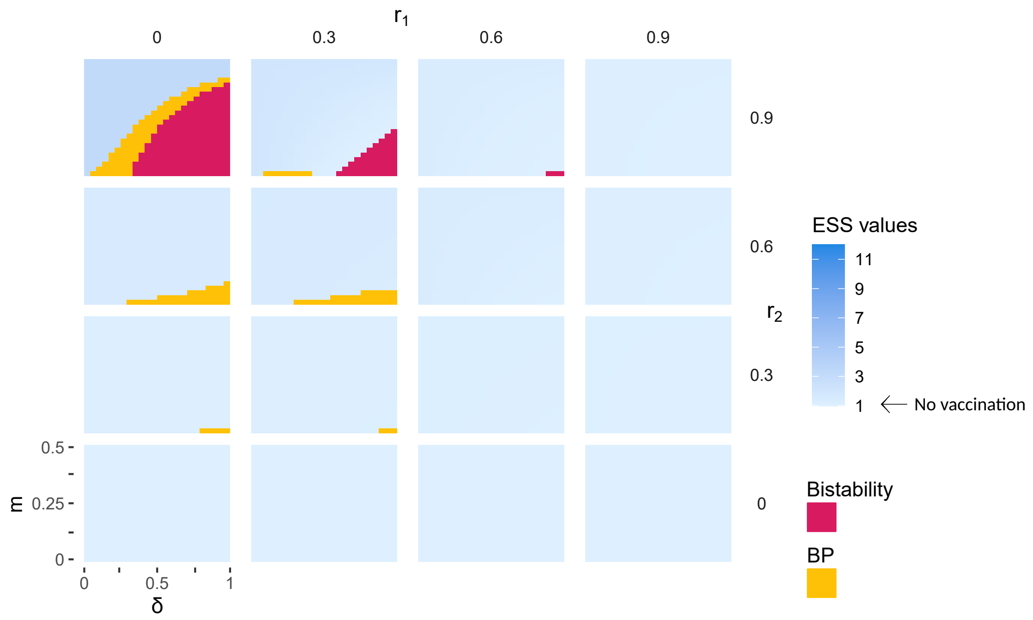


**(b) Anti-growth ($r_2$) with anti-transmission ($r_3$)**

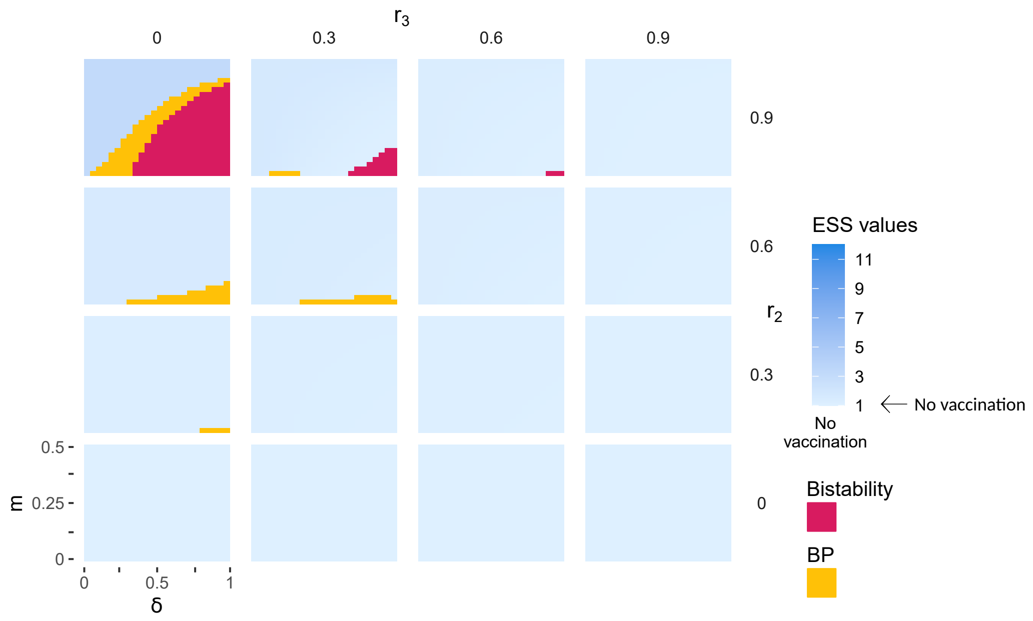


**Figure S.5:** Long-term evolutionary state calculated using adaptive dynamics, for anti-growth vaccine combined with (a) anti-infection and (b) anti-transmission, as a function of to $m$ and $\delta$. The region with a branching point (BP) is represented in yellow, the bistability region in pink, and the ESS region in a gradient of blue that represents the absolute value of ES virulence. Other parameters as in **figure 1** in the main text.

**(a) Anti-virulence ($r_4$) with anti-infection ($r_1$)**

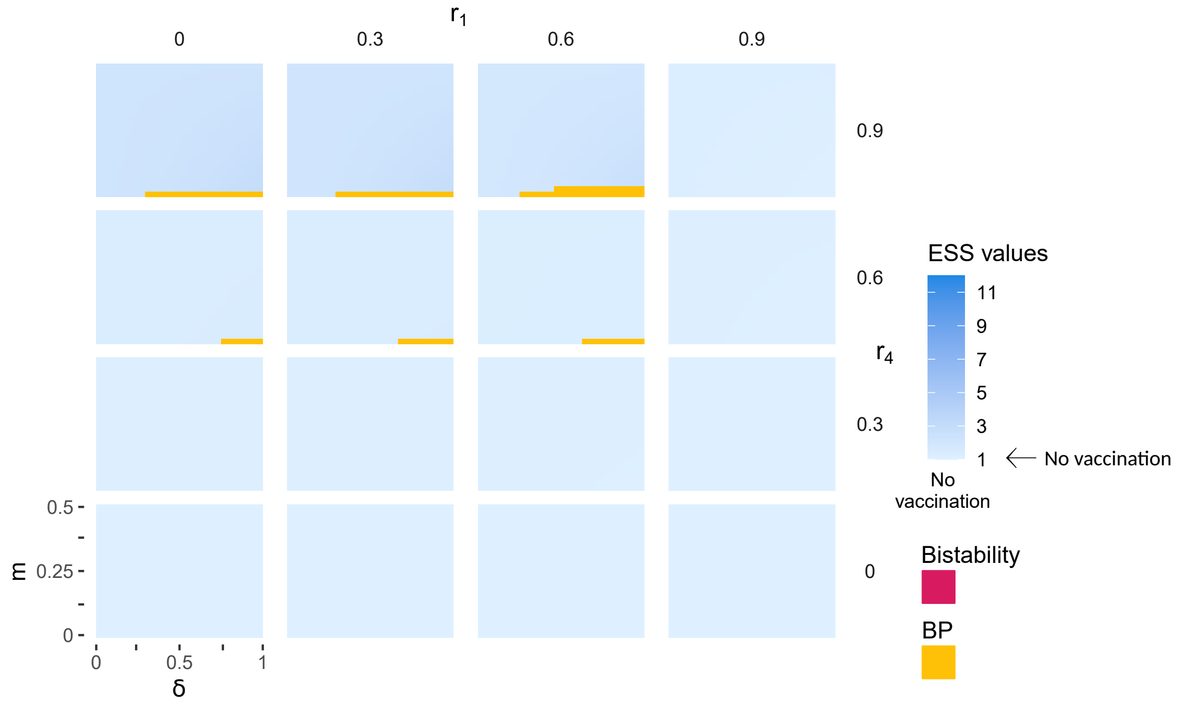


**(b) Anti-virulence ($r_4$) with anti-transmission ($r_3$)**

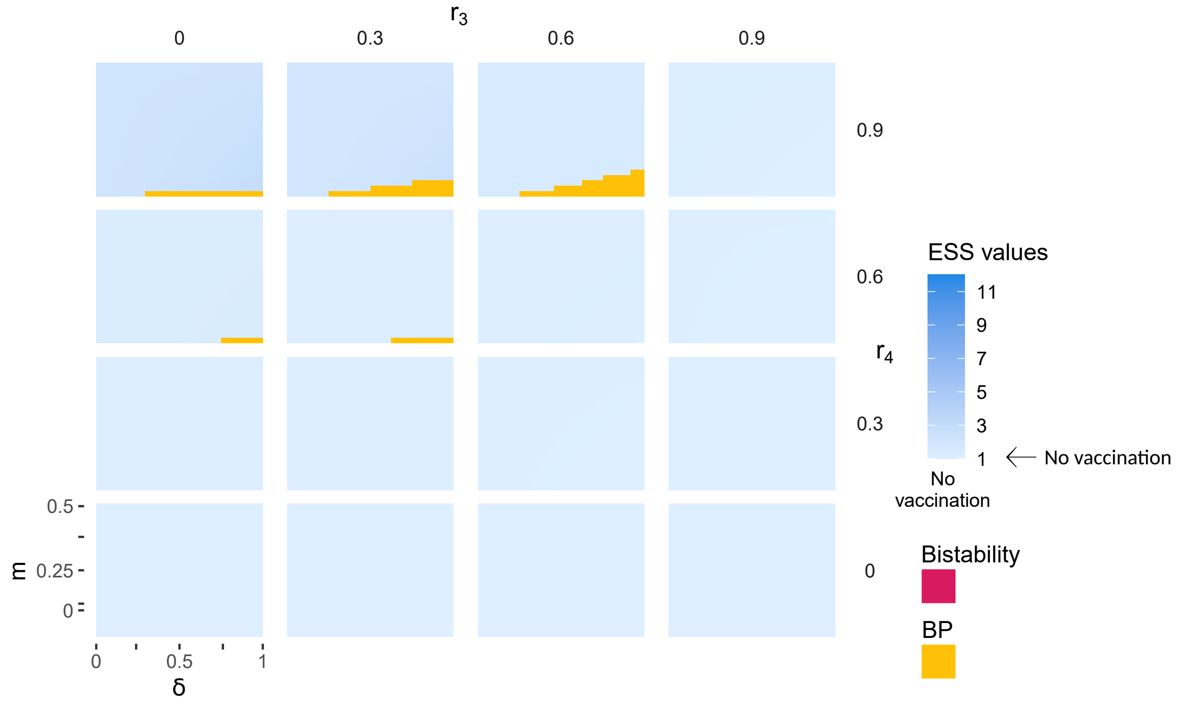


**Figure S.6:** Long term evolutionary state calculated using adaptive dynamics, for anti-virulence vaccine combined with (a) anti-infection and (b) anti-transmission, according to $m$ and $\delta$. See figure S.5 for the legend.

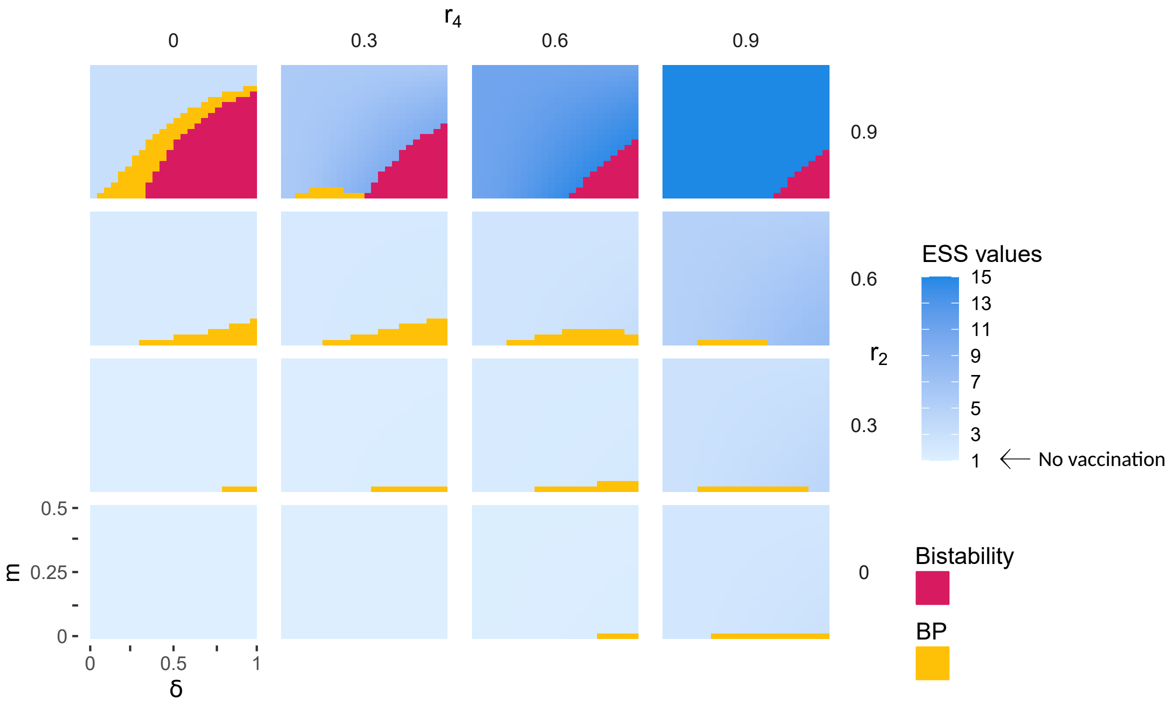


**Figure S.7:** Long-term evolutionary state calculated using adaptive dynamics, for anti-growth vaccine ($r_2$) combined with anti-virulence ($r_4$), as a function of $m$ and $\delta$. See figure S.5 for the legend.

### S.4 Evolution in a polymorphic population

In addition to our monomorphic adaptive dynamics anlaysis, we also numerically investigate a metapopulation model with a phenotypic distribution of pathogen strains. This polymorphic pathogen population evolves by mutation-selection until an equilibrium distribution is reached, which can be either unimodal or bimodal. We can thus compare the results with the AD predictions. Model eq. 1 in the main text is then modified into the following diffusion model :

$$\frac{\mathrm{d}S_A^N}{\mathrm{d}t} = (1-\nu_A)b - S_A^N(d + (1-m)h_{A,i} + mh_{B,i}),$$

$$\frac{\mathrm{d}S_A^T}{\mathrm{d}t} = \nu_A b - S_A^T(d + \sigma((1-m)h_{A,i} + mh_{B,i})), \qquad \text{(S.23)}$$

$$\frac{\mathrm{d}S_B^N}{\mathrm{d}t} = (1-\nu_B)b - S_B^N(d + (1-m)h_{B,i} + mh_{A,i}),$$

$$\frac{\mathrm{d}S_B^T}{\mathrm{d}t} = \nu_B b - S_B^T(d + \sigma((1-m)h_{B,i} + mh_{B,i})), \qquad \text{(S.24)}$$

$$\frac{\mathrm{d}I_{A,i}^N}{\mathrm{d}t} = S_A^N((1-m)h_{A,i} + mh_{B,i}) - (d + \alpha_A^N)I_{A,i}^T + \frac{\mathrm{Vm}}{2}\frac{\partial^2 I_{A,i}^N}{\partial z^2},$$

$$\frac{\mathrm{d}I_{A,i}^T}{\mathrm{d}t} = \sigma S_A^T((1-m)h_{A,i} + mh_{B,i}) - (d + \alpha_A^T)I_{A,i}^T + \frac{\mathrm{Vm}}{2}\frac{\partial^2 I_{A,i}^T +}{\partial z^2}$$

$$\frac{\mathrm{d}I_{B,i}^N}{\mathrm{d}t} = S_B^N((1-m)h_{B,i} + mh_{A,i}) - (d + \alpha_B^N)I_{B,i}^T + \frac{\mathrm{Vm}}{2}\frac{\partial^2 I_{B,i}^N}{\partial z^2},$$

$$\frac{\mathrm{d}I_{B,i}^T}{\mathrm{d}t} = \sigma S_B^T((1-m)h_{B,i} + mh_{A,i}) - (d + \alpha_B^T)I_{B,i}^T + \frac{\mathrm{Vm}}{2}\frac{\partial^2 I_{B,i}^T +}{\partial z^2},$$

where $i$ is the strain index which varies between 1 and $n$, and Vm is the variance of the mutation times the probability of the mutation (Débarre et al., 2013). We note that the susceptible equations remain very similar, but we have added a diffusion parameter in the infected hosts equations. Moreover, we have $4 \times n$ type of infected hosts. In numerical simulations, the diffusion parameter is discretised in class $k$

($N$ or $T$) as,

$$\frac{\partial^2 I_{A,i}^k}{\partial z^2} = \begin{cases} i \neq 1,\, i \neq n\,, & \frac{1}{\mathrm{d}z}\left(I_{A,i-1}^k - 2I_{A,i}^k + I_{A,i+1}^k\right) \\ i = 1\,, & \frac{1}{\mathrm{d}z}\left(I_{A,2}^k - I_{A,1}^k\right) \\ i = n\,, & \frac{1}{\mathrm{d}z}\left(I_{A,n-1}^k - I_{A,n}^k\right) \end{cases}$$

with d$z$ the mutational step.

The initial condition of the simulation is a Gaussian distribution for strains densities, which is typically centered on $z = 5$ with a variance of 1, and the strains values varies in 0.8 and 12 by 0.1.

The forces of infection becomes:

$$h_{j,i} = \sum_{i=1}^{n} \left(\beta_{j,i}^N I_{j,i}^N + \beta_{j,i}^T I_{j,i}^T\right) \tag{S.25}$$

### S.5 Numerical exploration of all combinations of $\nu_A$ and $\nu_B$, with different migration rates or efficacy, for anti-growth and anti-virulence vaccines

In the main text, we have restricted the study to the case $\nu = 0.5$, with $\nu_A \in [0; 0.5]$ and $\nu_B \in [0.5; 1]$. Here, we explore different values of average coverage, with $\nu_A \in [0; 1]$ and $\nu_B \in [0; 1]$ with anti-virulence or anti-growth vaccine, with different vaccine efficacy, for different value of pathogen migration ($m$).

**(a) Anti-virulence vaccine** ($r_4 = 0.7$)

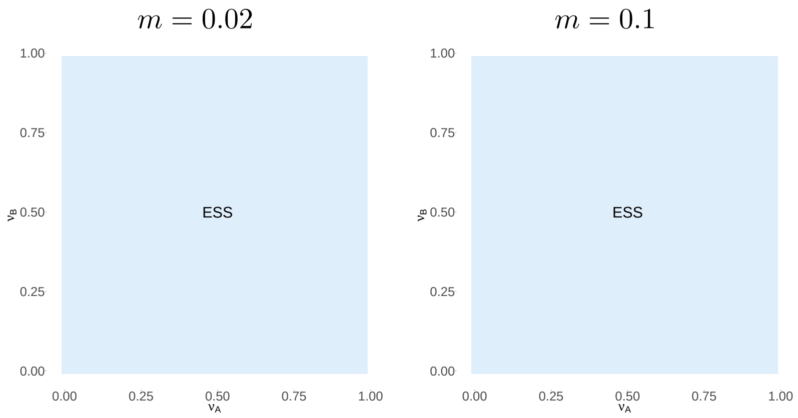


**(b) Anti-virulence vaccine** ($r_4 = 0.9$)

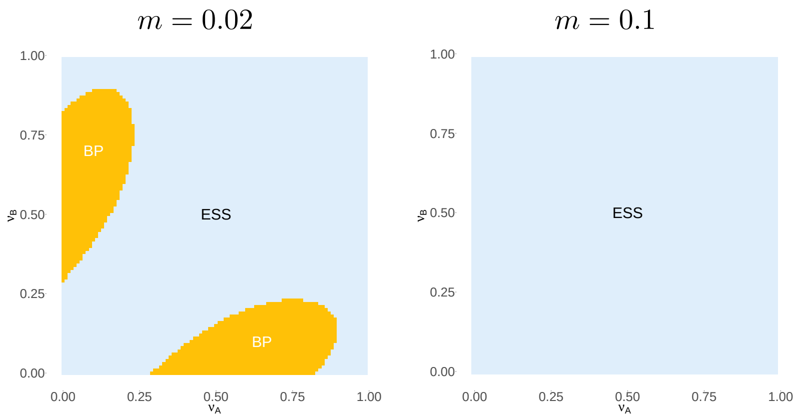


**Figure S.8:** Long term evolutionary state calculated using adaptive dynamics, for anti-virulence vaccines, with (a) $r_4 = 0.7$ and (b) $r_4 = 0.9$ with $m = 0.02$ or $m = 0.1$. ESS are represented in blue, branching point (BP) in yellow and bistability in pink.

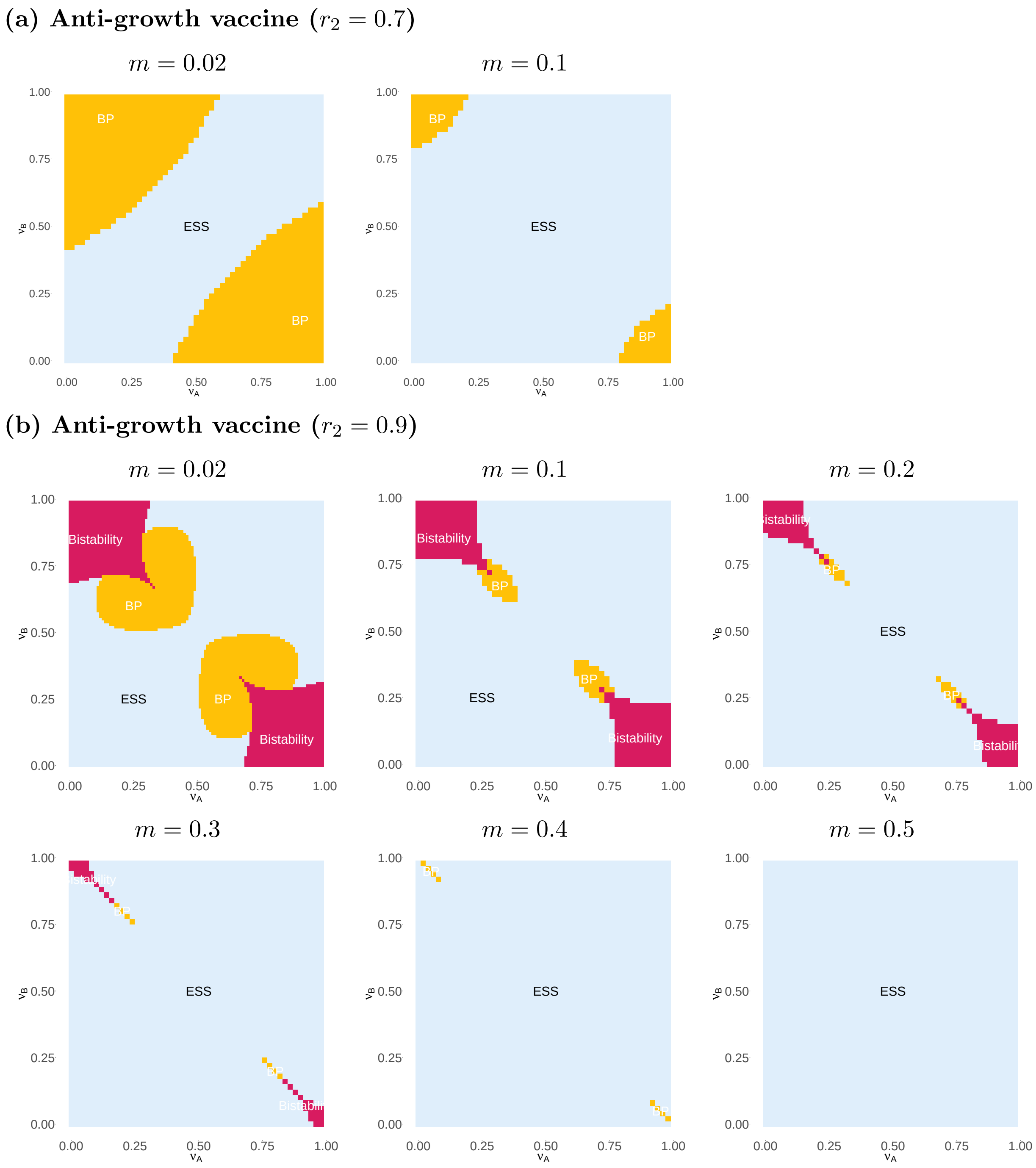


**Figure S.9:** Long term evolutionary state calculated using adaptive dynamics, for anti-growth vaccines, with (a) $r_2 = 0.7$ and $m = 0.02$ or $m = 0.1$ and (b) $r_2 = 0.9$ $m = 0.02\,,0.1\,,0.2\,,0.3\,,0.4\,,0.5$. ESS are represented in blue, branching point (BP) in yellow and bistability in pink.

### S.6 Prevalence and death flow

Here, we highlight that the anti-virulence vaccine increases significantly the prevalence and the death flow, regardless of the numbers of strains that coexists. The same observation applies to the anti-growth vaccine, although in smaller proportions. Thus, polymorphism affects neither prevalence nor the cumulative number of deaths, although, as we note in the main text, this will depend on the exact assumptions we make on the epidemiological dynamics.

The prevalence in the metapopulation is numerically calculated at the singularity, as $\sum_k \sum_i = I_i^k/(S_i^k + I_i^k)$, with $i$ the class ($N$ or $T$) and $k$ the population ($A$ or $B$). The death flow, $\overline{\alpha}$ is numerically calculated in a polymorphic population of pathogens, whose traits varies between 0 and 11 by 0.1 (111 strains). The strains are initially present in equivalent frequencies, and we calculate the death flow at equilibrium as $\overline{\alpha} = \sum_i \sum_k \alpha_i^k I_i^k/(S_i^k + I_i^k)$.

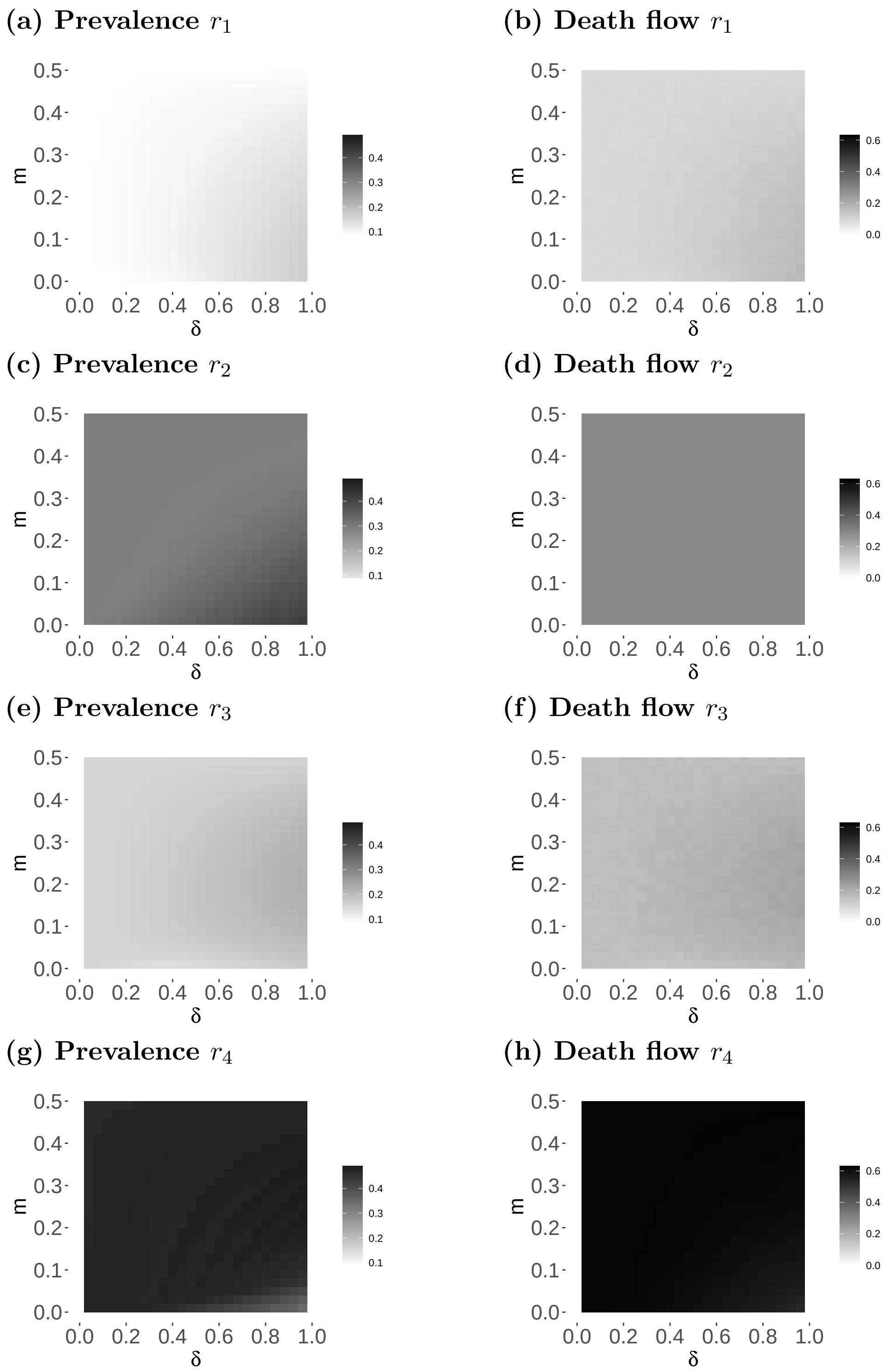


**Figure S.10:** The prevalence in the metapopulation numerically calculated at the singularity, as $\sum I_i^k/(S_i^k + I_i^k)$, with $i$ the class ($N$ or $T$) and $k$ the population ($A$ or $B$) and the death flow numerically calculated in a polymorphic population of pathogens, whose traits varies between 0 and 11 by 0.1 (111 strains). The strains are initially present in equivalent frequencies ($10^{-4}$), and we calculate the death flow at equilibrium as $\overline{\alpha} = \sum_i \sum_k \alpha_i^k I_i^k/(S_i^k + I_i^k)$, for (a, b) anti-infection, (c, d) anti-growth, (e, f) anti-transmission and (g, h) anti-virulence vaccine, with $r_i = 0.9$, and the other parameters as in **figure 1** in the main text.

### S.7 Adult vaccination

In the main text, the model implicitly considers that only newborn hosts are vaccinated. Here, we modify the model to account for vaccination of adult hosts and present additional results showing that the main qualitative results of the model in the main text are preserved. With adult vaccination, the dynamics become

$$\begin{aligned}
\frac{\mathrm{d}S_A^N}{\mathrm{d}t} &= b - \nu_A S_A^N + S_A^N(d + (1-m)h_A + mh_B)\,, \\
\frac{\mathrm{d}S_A^T}{\mathrm{d}t} &= \nu_A S_A^N - S_A^T(d + \sigma((1-m)h_A + mh_B))\,, \\
\frac{\mathrm{d}S_B^N}{\mathrm{d}t} &= b - \nu_B S_B^N + S_B^N(d + (1-m)h_B + mh_A)\,, \\
\frac{\mathrm{d}S_B^T}{\mathrm{d}t} &= \nu_B S_B^N - S_B^T(d + \sigma((1-m)h_B + mh_A))\,, \\
\frac{\mathrm{d}I_A^N}{\mathrm{d}t} &= S_A^N((1-m)h_A + mh_B) - (d + \alpha_A^N)I_A^N\,, \\
\frac{\mathrm{d}I_A^T}{\mathrm{d}t} &= \sigma S_A^T((1-m)h_A + mh_B) - (d + \alpha_A^T)I_A^T\,, \\
\frac{\mathrm{d}I_B^N}{\mathrm{d}t} &= S_B^N((1-m)h_B + mh_A) - (d + \alpha_B^N)I_B^N\,, \\
\frac{\mathrm{d}I_B^T}{\mathrm{d}t} &= \sigma S_B^T((1-m)h_B + mh_A) - (d + \alpha_B^T)I_B^T\,,
\end{aligned} \tag{S.26}$$

with $b$ the birth rate, $d$ the natural death rate, $m$ the migration between the two subpopulations, that varies between 0 (no migration) and 0.5 (full migration), and $\nu_A$ and $\nu_B$ the vaccination rates in the supopulations $A$ and $B$. Note that, in the adult vaccination model, we are dealing with rates, which vary between $[0; \infty[$, and not with probabilities.

In **figure S.11** and **S.12**, we present the long term stability of the virulence evolution according to $\nu_A$ and $\nu_B$. **Figure S.13** can be compared with to the **figure 3** in the main text.

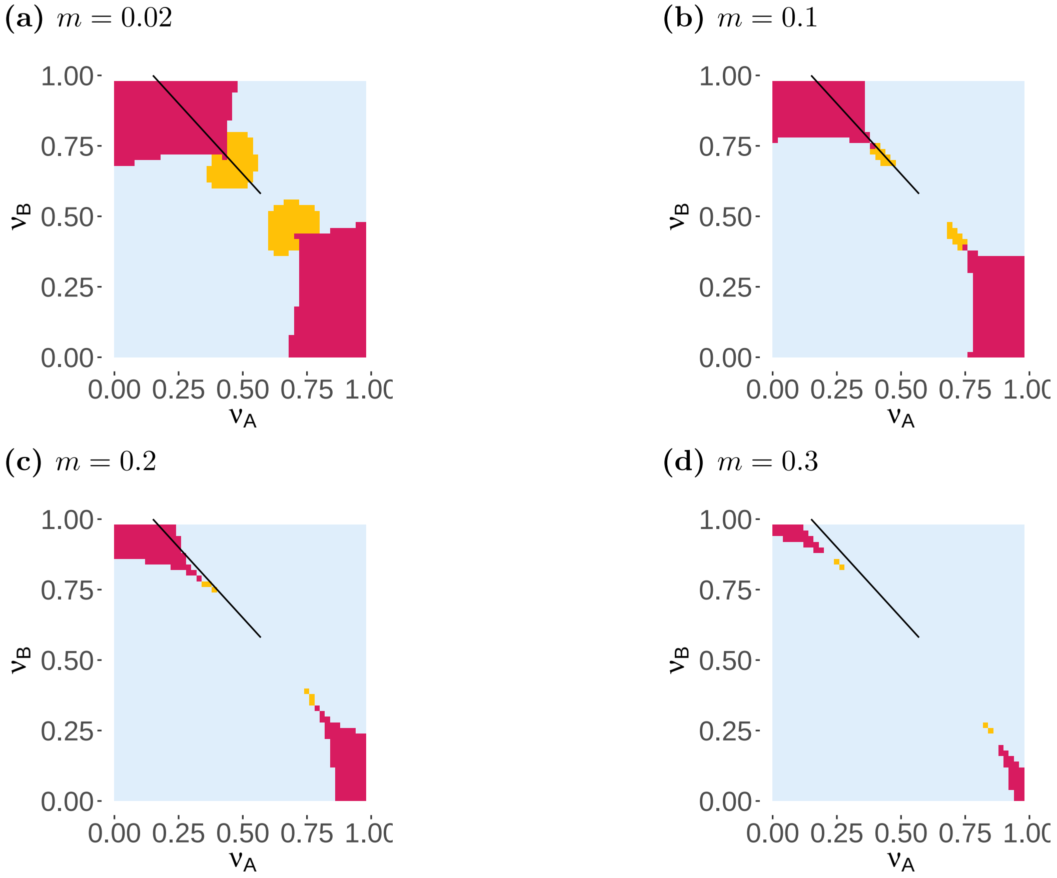


**Figure S.11:** Anti-growth ($r_2$). Long-term evolutionary state calculated using AD, according to the coverage in the subpopulations $A$, $f(\nu_A)$ and $B$, $f(\nu_B)$, where $f(\nu) = (1-\nu)/\nu$, for different values of migration, $m$. ESS's are represented in blue and branching points in yellow. The black lines represent the values of $\nu_A$ and $\nu_B$ used to calculate $\delta$ in the **figure S.13**. All produced with $b = 1.5$, $d = 1$, $r_2 = 0.9$.

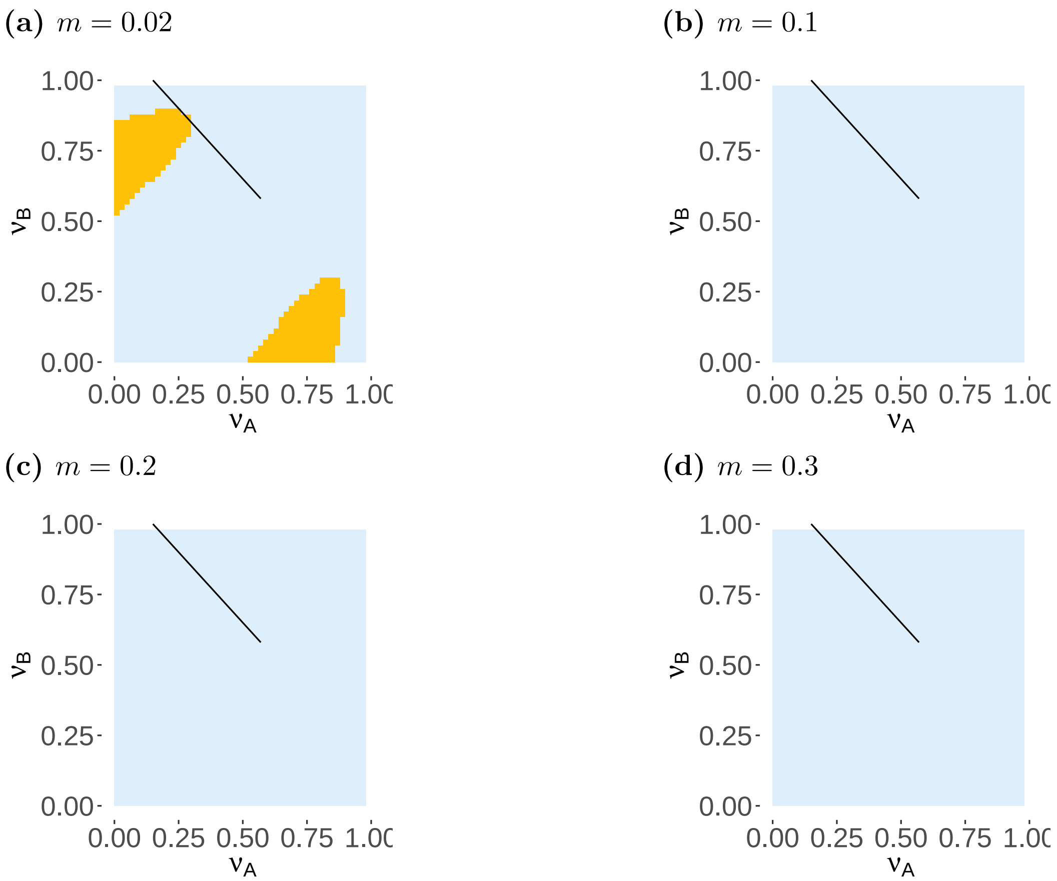


**Figure S.12:** Anti-virulence ($r_4$). Long-term evolutionary state calculated using AD, according to the coverage in the subpopulations $A$, $f(\nu_A)$ and $B$, $f(\nu_B)$, where $f(\nu) = (1-\nu)/\nu$, for different values of migration, $m$. ESS's are represented in blue and branching points in yellow. The black lines represent the values of $\nu_A$ and $\nu_B$ used to calculate $\delta$ in the **figure S.13**. All produced with $b = 1.5$, $d = 1$, $r_4 = 0.9$.

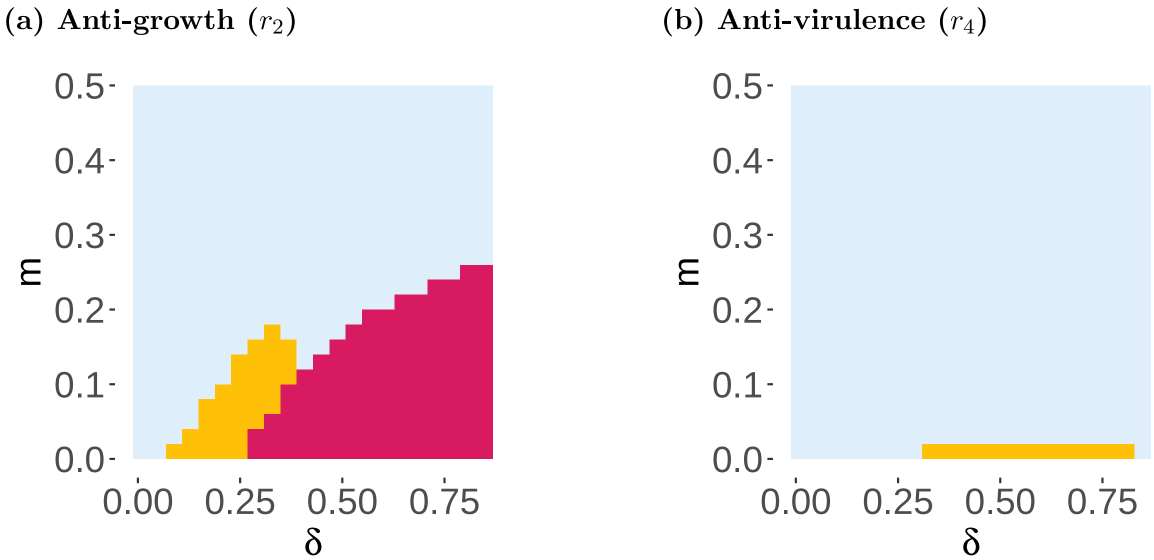


**Figure S.13:** Long-term evolutionary state calculated using adaptive dynamics, for (a) anti-growth ($r_2$) and (b) anti-virulence ($r_4$), ESS are represented in blue, BP in yellow and bistability in pink, according to $m$ and $\delta$, with $\nu = 0.575$ and $\delta = \nu_B - \nu_A$.
